# Dynamics and Evolutionary Conservation of B Complex Protein Recruitment During Spliceosome Activation

**DOI:** 10.1101/2024.08.08.606642

**Authors:** Xingyang Fu, Aaron A. Hoskins

## Abstract

Spliceosome assembly and catalytic site formation (called activation) involve dozens of protein and snRNA binding and unbinding events. The B-complex specific proteins Prp38, Snu23, and Spp381 have critical roles in stabilizing the spliceosome during conformational changes essential for activation. While these proteins are conserved, different mechanisms have been proposed for their recruitment to spliceosomes. To visualize recruitment directly, we used Colocalization Single Molecule Spectroscopy (CoSMoS) to study the dynamics of Prp38, Snu23, and Spp381 during splicing in real time. These proteins bind to and release from spliceosomes simultaneously and are likely associated with one another. We designate the complex of Prp38, Snu23, and Spp381 as the B Complex Protein (BCP) subcomplex. Under splicing conditions, the BCP associates with pre-mRNA after tri-snRNP binding. BCP release predominantly occurs after U4 snRNP dissociation and after NineTeen Complex (NTC) association. Under low concentrations of ATP, the BCP pre-associates with the tri-snRNP resulting in their simultaneous binding to pre-mRNA. Together, our results reveal that the BCP recruitment pathway to the spliceosome is conserved between *S. cerevisiae* and humans. Binding of the BCP to the tri-snRNP when ATP is limiting may result in formation of unproductive complexes that could be used to regulate splicing.

**KEY POINTS:**

- Prp38, Snu23, and Spp381 associate together to form the B Complex Proteins (BCP) Complex
- During yeast spliceosome assembly, the BCP binds after the tri-snRNP and leaves after NTC arrival
- At low ATP, the BCP pre-associates with the tri-snRNP in complexes that are likely unproductive

## INTRODUCTION

RNA splicing is the process of removing introns from precursor mRNAs (pre-mRNAs) concomitant with ligation of the flanking exons. The process is mediated by a megadalton ribonucleoprotein complex, the spliceosome. The components of spliceosomes can be broadly categorized as small nuclear ribonucleoprotein complexes (U1, U2, U4, U5, U6 snRNPs), protein-only complexes (Nineteen Complex, NTC; NTC-related complexes, NTR), and transiently interacting protein splicing factors (*e.g.*, several DExD/H-box ATPases required for splicing). While spliceosome assembly is largely ordered, it is also a highly dynamic process (**Fig. 1A**) (1–3). In brief, spliceosome assembly begins by recruitment of the U1, U2, and U4/U6.U5 tri-snRNPs to pre-mRNA to form the pre-B complex spliceosome. This is then followed by spliceosome activation during which the nascent active site is formed, the U1 and U4 snRNPs are released, and the NTC and NTR complexes are recruited. These steps involve the sequential formation of the B and B^act^ intermediate complexes. Ultimately, large-scale conformational and compositional changes lead to the formation of a U2/U6 di-snRNA catalytic center associated with the intron and capable of catalyzing splicing chemistry. Upon completion of exon ligation, the spliceosome disassembles, and its components are recycled for processing of other pre-mRNAs.

**Figure 1.**
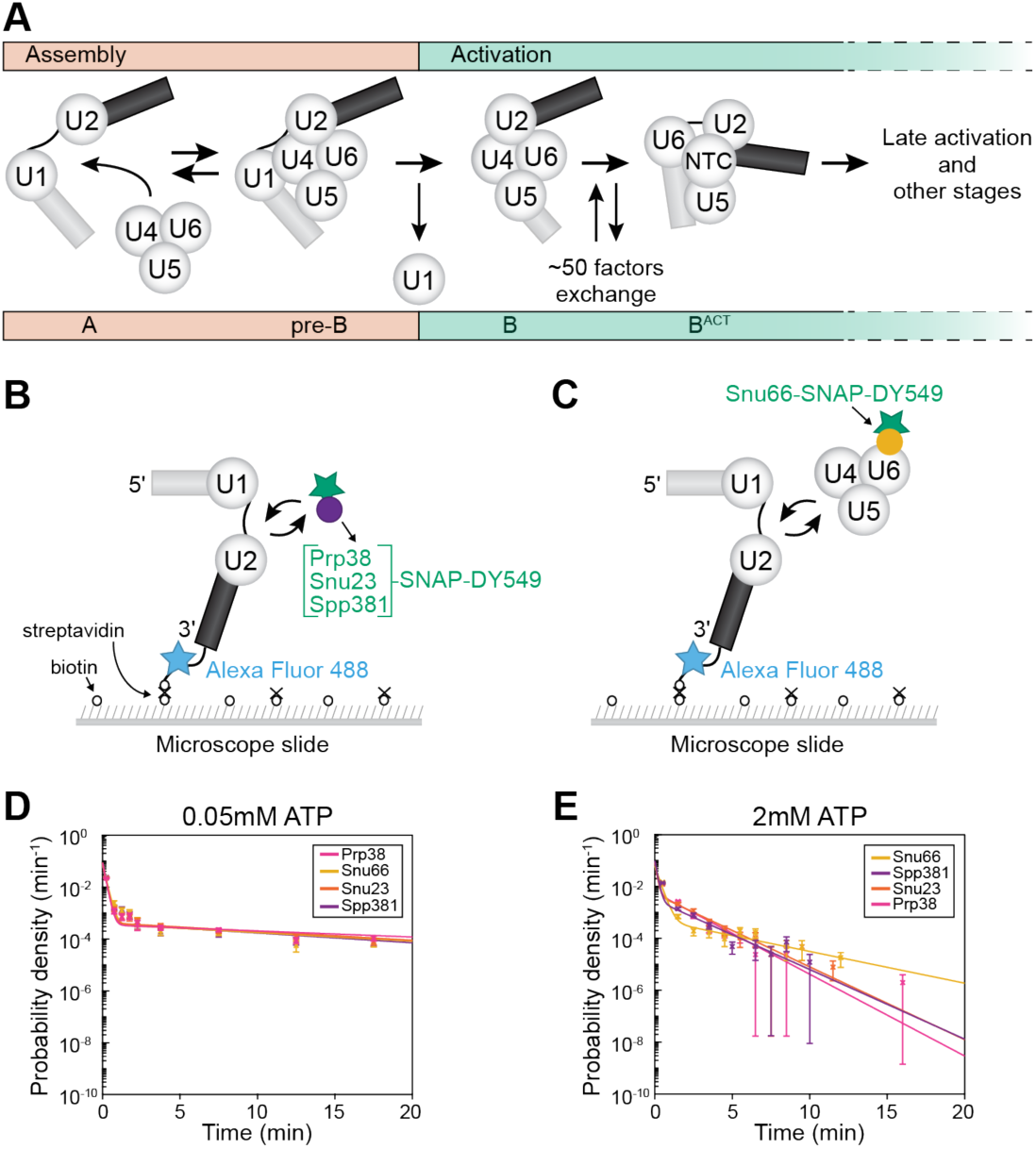
Overview of yeast spliceosome assembly and activation and CoSMoS assays. (**A**) Spliceosome activation begins after the tri-snRNP joins the spliceosome A complex to form the pre-B complex. Release of the U1 snRNP is then followed by rearrangements that include exchange of ∼50 protein and snRNA factors to form the activated spliceosome (B^act^). This panel was adapted from Fu *et al.* (19). (**B, C**) Schematics of 2-color CoSMoS experiments for observing either Prp38, Snu23, or Spp381 binding dynamics (Panel B) or Snu66 binding dynamics (Panel C). (**D, E**) Probability density histograms for dwell times of Prp38, Snu66, Snu23, or Spp381 under conditions that either block (0.05 mM ATP, Panel D) or permit spliceosome activation (2 mM ATP, Panel E).

During the activation of human spliceosomes, B complex-specific factors (PRPF38, MFAP1, ZMAT2, FBP21, SMU1, RED, NPW38, NPW38BP) are recruited during the transition from the pre-B to B complex spliceosome and released during the transition from the B to B^act^ complex (4–7). Cryo-EM structures of human spliceosomes have revealed where these factors are bound. ZMAT2 and PRPF38 are positioned to anchor the U6 snRNA/5’ splice site (SS) duplex to the N-terminal domain of PRP8 whereas MFAP1 helps to stabilize the interaction between the 5’ exon and U5 snRNA and interacts with the PRP8 switch loop (5). The binding sites for these proteins are mutually exclusive with that of Prp28 (a DEAD-box ATPase involved in the transfer of the 5’ SS from the U1 to the U6 snRNA) indicating that Prp28 must be released before they can bind at that location (8). Consequently, presence of these proteins on the human spliceosome signifies a particular state in which activation has begun, the pre-B complex has been converted to the B complex, the U1 snRNP has dissociated, the 5’ SS has been identified and transferred to U6, and Prp28 has departed.

Only PRPF38, ZMAT2 and MFAP1 are evolutionarily conserved between human and *S. cerevisiae* (hereafter yeast; Prp38, Snu23 and Spp381, respectively) (9). However, the three yeast proteins can be isolated as part of the yeast U4/U6.U5 tri-snRNP in the absence of splicing (9–13). As a result, two different pathways for recruitment of B-specific factors have been proposed: ordered recruitment of the human proteins during spliceosome activation after association of the tri-RNP (5) or simultaneous recruitment of the yeast tri-snRNP and B-specific factors (14). Given the evolutionarily conservation of the splicing machinery, the origins and consequences for this variation remains elusive as do the implications for 5’ SS identification and Prp28 function (9).

Prp38, Snu23 and Spp381 are all essential for yeast viability (15–17). Though no crystal structures of a yeast Prp38-Snu23-Spp381 co-complex have yet been solved, a crystal structure containing a ternary complex formed from fragments of *Chaetomium thermophilum* Prp38, Snu23, and MFAP1 has been obtained (18). This structure and accompanying biochemical data suggest that they can potentially function as a heterotrimeric subcomplex during splicing and associate with one another in the absence of the spliceosome. Further, the yeast proteins can form an isolable complex by gel filtration experiments (9). Despite evidence for interactions between the three proteins, it is not known if either the yeast or human B-specific factors are recruited to and release from tri-snRNPs or spliceosomes individually or as a single subunit.

Since the transition from B to B^act^ spliceosome involves the exchange of about fifty factors (including the release of BCPs and the U4 snRNP and recruitment of the NTC and NTR), it is unsurprising that this transition likely involves the formation of several intermediates (14,19). Two recent human pre-B^act^ spliceosome structures stalled during this transition using a small molecule splicing inhibitor revealed at least two sub-steps (5). In pre-B^act-1^ spliceosome (representing the first sub-step), U4 snRNP has been released, PRPF38, ZMAT2 and MFAP1 are present, and a subset of NTC proteins have associated and can be resolved in the structure. PRPF38, ZMAT2 and MFAP1 are also retained in pre-B^act-2^ where they function to impede docking of the U6/5’ SS duplex into the nascent active site and Prp8 re-arrangement. Together, these data suggest that PRPF38, ZMAT2 and MFAP1 are released after U4 snRNP dissociation and after NTC recruitment by an unknown mechanism. Similar activation intermediates have not yet been structurally characterized for the yeast spliceosome.

The highly compositional and conformational dynamic activation process renders the underlying mechanism intrinsically complicated and difficult to study. Even though cryo-EM structures have provided a few snapshots along the activation path, the number of kinetic steps and characteristic spliceosome complexes involved in each transition are challenging to predict. Previously, our lab has employed Colocalization Single Molecule Spectroscopy (CoSMoS) to study the dynamics of major factors (U4 snRNP proteins, NTC proteins and Lsm2-8 proteins) involved in the activation pathway (19,20). In this work, we analyze the dynamics of yeast Prp38, Snu23 and Spp381 during activation and compare these to other splicing factors to order their arrival to and departure from single spliceosomes. Our results show that a human-like activation pathway is conserved for the yeast splicing machinery. Prp38, Snu23 and Spp381 bind to and dissociate from spliceosomes as a subcomplex, which we refer to as the B Complex Protein (BCP) subcomplex. Under splicing conditions, spliceosomes assemble by ordered, sequential addition of the tri-snRNP and BCP to pre-mRNAs. Surprisingly, association of the BCP with the tri-snRNP is ATP-dependent. Under low ATP conditions that prevent splicing, the BCP and tri-snRNP instead associate with pre-mRNA simultaneously. This observation likely explains the presence of the yeast BCP in tri-snRNPs and tri-snRNP-containing complexes purified under non-splicing conditions. Together, our results provide insight into how tri-snRNP composition may be regulated by ATP and demonstrate evolutionary conservation of a fundamental step in spliceosome activation.

## MATERIALS AND METHODS

### Yeast Strains

Yeast strains (**Supplementary Table S1**) were derived from the protease-deficient strain BJ2168. Splicing factors were C-terminally tagged by integrating fast SNAP or DHFR tags at the appropriate genomic locations by homologous recombination as described (1,20).

### Yeast Growth Assay

Selected yeast strains were grown in YPD media to saturation in a 30°C shaking incubator. Concentrated yeast cultures were then diluted to OD_600_ = 0.03 with YPD media and placed in a 48-well plate (Corning Costar 48-well Flat Bottom Cell Culture Plate). The plate was covered with a Breathe-Easy plate sealing membrane to limit evaporation during incubation. A Tecan Infinite 200Pro plate reader was then used to monitor yeast growth by measuring the OD_600_ for 30 h at 30 °C with shaking.

### Preparation of Yeast Whole Cell Extract (WCE)

Yeast WCE for splicing was prepared according to published protocols using the ball mill method (21). SNAP-tagged proteins were fluorophore labeled as previously described (22). Briefly, SNAP-Surface® 549 dye (S9112S, New England BioLabs; abbreviated elsewhere as SNAP-DY549) in DMSO was added to 1.2 mL yWCE to a final concentration of 1 μM. The reaction tube was rotated in the dark for 30 min at room temperature. The reaction was then loaded onto a pre-equilibrated G-25 Sephadex column (Kontes Flex Column) in SEC buffer (25 mM HEPES-KOH pH 7.9, 50 mM KCl, 1 mM DTT, 10% v/v glycerol) at 4°C to remove excess dye. A peristaltic pump was used for pre-equilibrating the column and eluting the labeled extract at a flow rate of ∼0.25 mL/min with the SEC buffer. Fluorophore labeling of the proteins was confirmed by SDS-PAGE and fluorescence using a Typhoon FLA 9000 scanner (Cytiva) at 532nm. Results were analyzed with ImageQuantTL (Cytiva) software.

### In Vitro Splicing Assays

[α-^32^P] UTP radio-labeled (PerkinElmer) and m^7^G(5′)ppp(5′)G capped (New England Biolabs) RP51A pre-mRNA substrates were made by *in vitro* transcription of a linear DNA template with T7 RNA polymerase (Agilent or purified in the laboratory). The DNA template was produced from a PCR reaction of pBS117 plasmid (23) using Taq DNA polymerase (M7122, Promega), followed by gel purification of the products with a Wizard SV Gel and PCR Clean-Up System kit (Promega). Transcription products were separated on a 6% denaturing polyacrylamide gel (AccuGel 19:1, National Diagnostics), followed by ethanol precipitation of the extracted RP51A transcripts. Transcripts were resuspended in nuclease-free water (Ambion, Fischer Scientific) and quantified with a liquid scintillation counter (LSC, tri-carb 2900TR, Packard).

A typical *in vitro* splicing reaction included 40% v/v WCE and 0.2 nM RP51A substrate in a splicing buffer [final concentrations: 100mM KPi pH 7.3, 3% w/v PEG-8000, 1mM DTT, 2.5mM MgCl_2_, 0.2U/μL RNasin Plus (Promega)]. The reaction was incubated at room temperature for 45 min. The reaction was quenched in a splicing dilution buffer (100mM Tris base pH 7.5, 10mM EDTA pH 8.0, 1% w/v SDS, 150mM NaCl, 300 mM NaOAc pH 5.3). RNAs from the reaction were extracted using phenol-chloroform, ethanol precipitated, resuspended in deionized formamide, and separated on a 12% denaturing polyacrylamide gel. Gels were then dried and exposed to a phosphor screen overnight. The screen was imaged with a Typhoon FLA 9000 scanner (Cytiva), and results were analyzed with ImageQuantTL (Cytiva) software. The intensities efor RP51A pre-mRNAs and splicing products were determined by integrating the signals within same-sized rectangles around the band. The background corrected intensities for bands were then used for calculating splicing efficiencies. The 1^st^ step efficiency was calculated as the ratio of the sum of the intensities of bands corresponding to products having completed 5’ SS cleavage (the mRNA and intron lariat-3’ exon) over the sum of those intensities plus that for the pre-mRNA band. The 2^nd^ step efficiency was calculated as the ratio of the intensity of the mRNA band to the sum of the intensities of the pre-mRNA, intron lariat-3’ exon, and mRNA bands.

### Preparation of Fluorescently-Labeled RP51A pre-mRNAs

Fluorescent, biotinylated RP51A pre-mRNAs were prepared as described by splinted ligation of a fluorescent biotin handle oligonucleotide to a capped, RP51A transcript prepared by *in vitro* transcription using T4 RNA Ligase 2 (19,24).

### CoSMoS Assays

Microscope slides (100490-396, VWR) and cover glasses (12-553-455, Fischer) were cleaned and passivated using a mixture of mPEG-SVA (MPEG-SVA-5K, Laysan Bio) and mPEG-biotin-SVA (BIO-PEG-SVA-5K, Laysan Bio) at a ratio of 1:100 w/w in 100mM NaHCO_3_ (pH 8) buffer as previously described (19).

For each CoSMoS assay, the mPEG mixture was washed off the slide with 1x PBS and then streptavidin (0.2 mg/mL, SA10, Kelowna International Scientific) was added and allowed to bind the surface. The excess streptavidin was washed off the surface using PBS and the biotinylated, fluorescent RNAs were then bound. The slide was washed a final time with PBS before addition of splicing assay buffer containing 40% v/v WCE, 20 nM Cy5-TMP, oxygen scavenger (5 mM PCA and 0.96 U/mL PCD), 1 mM Trolox, and triplet quenchers (0.5 mM propyl gallate, 1 mM cyclooctatetraene, 1 mM 4-nitrobenzyl alcohol) was added (1). The triplet quenchers were added as a mixture from a 100x stock in DMSO. Additional details of these procedures have been recently described elsewhere (19).

### CoSMoS Data Acquisition

Single molecule data were collected on a custom-built objective-type TIRF microscope using GLIMPSE software (https://github.com/gelles-brandeis/Glimpse) as previously described (19). For the laser illumination, powers were set to the ranges of 1.2 – 1.5 mW, 500 – 600 μW, 400-440 μW, and 2.5 mW for the 488 nm (blue), 532 nm (green), 633 nm (red) and 785 nm (infrared) lasers, respectively. The 488, 532, and 633 nm lasers were used to image fluorophores while the 785 nm laser was used for focusing.

Cycles of time-lapse imaging were used according to the following excitation scheme with each frame lasting 1s: the sample was illuminated with the 785 nm for focusing, the 488 nm blue laser was then turned on to collect two consecutive frames to image the immobilized RNAs, and then the 532 and 633 nm lasers were turned on to simultaneously collect 14 frames with a 3 s delay between adjacent frames. The total cycle time was ∼1 min and this cycle was repeated 50x to collect videos lasting for ∼50 min. To avoid photo bleaching of DY-549 and Cy5 fluorophores by the 488 nm laser, the path of the laser was physically blocked after collection of 10 frames of blue laser-illuminated images.

### Photobleaching Control

To estimate the influence of photobleaching of the SNAP tag fluorophore (DY549) on the experiments, we utilized a purified, biotinylated, and fluorescently-labeled SNAP protein. The protein was imaged under the same conditions used to follow splicing reactions, and its average fluorophore lifetime (τ = 995 s) was determined by measuring the loss in fluorescence signal overtime as previously described (19).

### CoSMoS Data Analysis

Single molecule data were analyzed as described previously (1,19,25) using a custom program imscroll (https://github.com/gelles-brandeis/CoSMoS_Analysis) written in MATLAB (The Mathworks). Binding events were identified as signals centered within the AOI that appeared with intensities greater than 3.6x the standard deviation above the mean of the background noise. Loss of signals were identified as points in time at which the signal fell below 1x standard deviation above the mean background.

Analysis of the measured dwell times and fits to kinetic equations were carried out using MATLAB and AGATHA software (https://github.com/hoskinslab/AGATHA) using maximum likelihood methods and fitting to equations containing one (Eq. 1), two (Eq. 2), or three (Eq. 3) exponential terms as described (25).

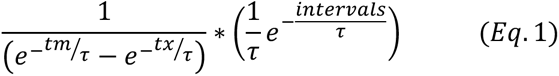

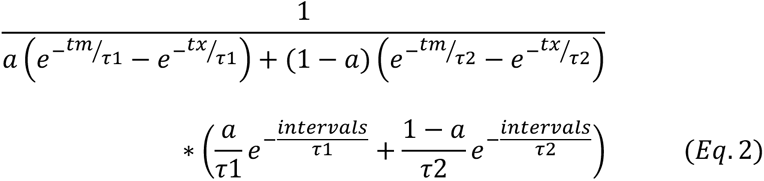

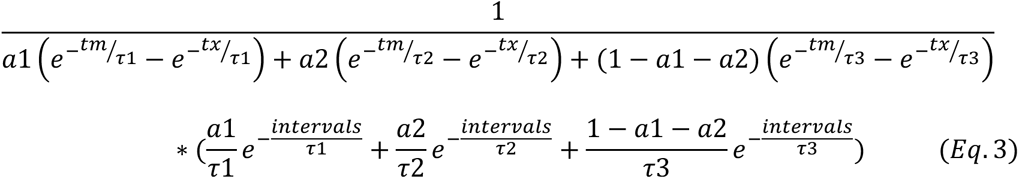

Bootstrapping was used to calculate standard errors for all fitted parameters. Fit parameters are included in **Supplementary Tables S2** and **S3**. Histograms for the distribution of events were generated in MATLAB with empty bins removed. The error (standard deviation) for each bin were calculated using Eq. 4, with the assumption that the number of events within a bin follows a binomial distribution.

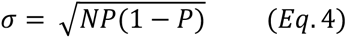

## RESULTS

### Prp38, Snu23 and Spp38 Have Similar pre-mRNA Binding Kinetics

To visualize splicing factor dynamics directly, we genetically encoded C-terminal fast SNAP tags on Prp38, Snu23, and Spp381 in three separate yeast strains (**Supplementary Table S1**, strains #5-7). As a control, we also labeled the tri-snRNP protein Snu66 in a separate strain (**Supplementary Table S1**, strain #4). Snu66 is released during the B to B^act^ transition but occupies a different binding site on the spliceosome than the other three factors (12). We confirmed that these SNAP-tagged proteins can be readily labeled with benzylguanine-fluorophores (SNAP-DY549) in yeast whole cell extracts (WCEs) and that the labeled extracts efficiently spliced RP51A substrate pre-mRNAs (**Supplementary Figs. S1, S2**). We do note, however, that strains containing tagged Prp38 grew more slowly and produced WCE with lower splicing activity when Snu23 and Spp381 or NTC components were also DHFR tagged (**Supplementary Fig. S2**) relative to others. It is possible that tagging of Prp38 disrupts expression of its neighboring gene, SMD1 (a component of the U snRNP Sm ring), and that this contributes to the observed differences in growth and splicing activity (26).

To observe splicing factor dynamics, we flowed a pre-mixed solution consisting of a fluorescently labeled WCE and splicing buffer into a reaction chamber containing surface-immobilized, Alexa Fluor 488-labeled RP51A pre-mRNAs (**Fig. 1B, C**). We carried out the experiments at two ATP concentrations that either permit spliceosome activation and splicing (2mM ATP) or inhibit activation and splicing but permit assembly (0.05mM ATP)(27). Using a custom-built CoSMoS microscope(28), we could observe binding and unbinding dynamics of fluorescently labeled proteins on the fixed fluorescent pre-mRNA molecules in real time.

All of the proteins showed ATP-dependent dynamics. At 0.05mM ATP, the dwell times of each protein bound to single pre-mRNA molecules often lasted for tens of minutes (**Supplementary Fig. S3**). In comparison, the binding events at 2 mM ATP were much shorter and lasted from a few seconds to a few minutes. These results resemble single molecule data obtained from other proteins involved in spliceosome activation, such as Prp3 and Prp4 from U4 snRNP complexes (20) and Lsm8 from the Lsm2-8 ring complex (19). It is likely that the long dwell times observed at 0.05 mM ATP originated from stalled spliceosome complexes while the shorter times seen at 2 mM ATP originated from transient binding due to release of these factors during spliceosome activation.

We further analyzed the binding dynamics by fitting the collected, unbinned dwell times to exponential-based functions (**Supplementary Table S2**) and generating probability density histograms. At low ATP, histograms and fits for Prp38, Spp381, Snu23, and Snu66 are very similar to one another (**Fig. 1D**, **Supplementary Table S2**). In contrast, under conditions that permit activation (2mM ATP), the distributions and fits for Prp38, Spp381 and Snu23 remained similar to one another but were distinct from those of Snu66 (**Fig. 1E, Supplementary Table S2**). Together, these results indicate that Prp38, Spp381, Snu23 interact with the splicing machinery with similar kinetic features. These features are shared with Snu66 only under low ATP conditions that inhibit spliceosome activation and splicing.

### Prp38, Spp381, Snu23 are Recruited to and Released from Spliceosomes Simultaneously

To see if the similarity in Prp38, Spp381, and Snu23 kinetics is coincidental or results from their presence together in a single spliceosomal subcomplex, we carried out experiments to visualize the recruitment and release of the proteins relative to one another during splicing. We carried out 3-color CoSMoS assays using WCE containing Prp38-SNAP labeled with SNAP-DY549 fluorophores and DHFR tags on Spp381 and Snu23 labeled with Cy5-TMP fluorophores (**Fig. 2A**).

**Figure 2.**
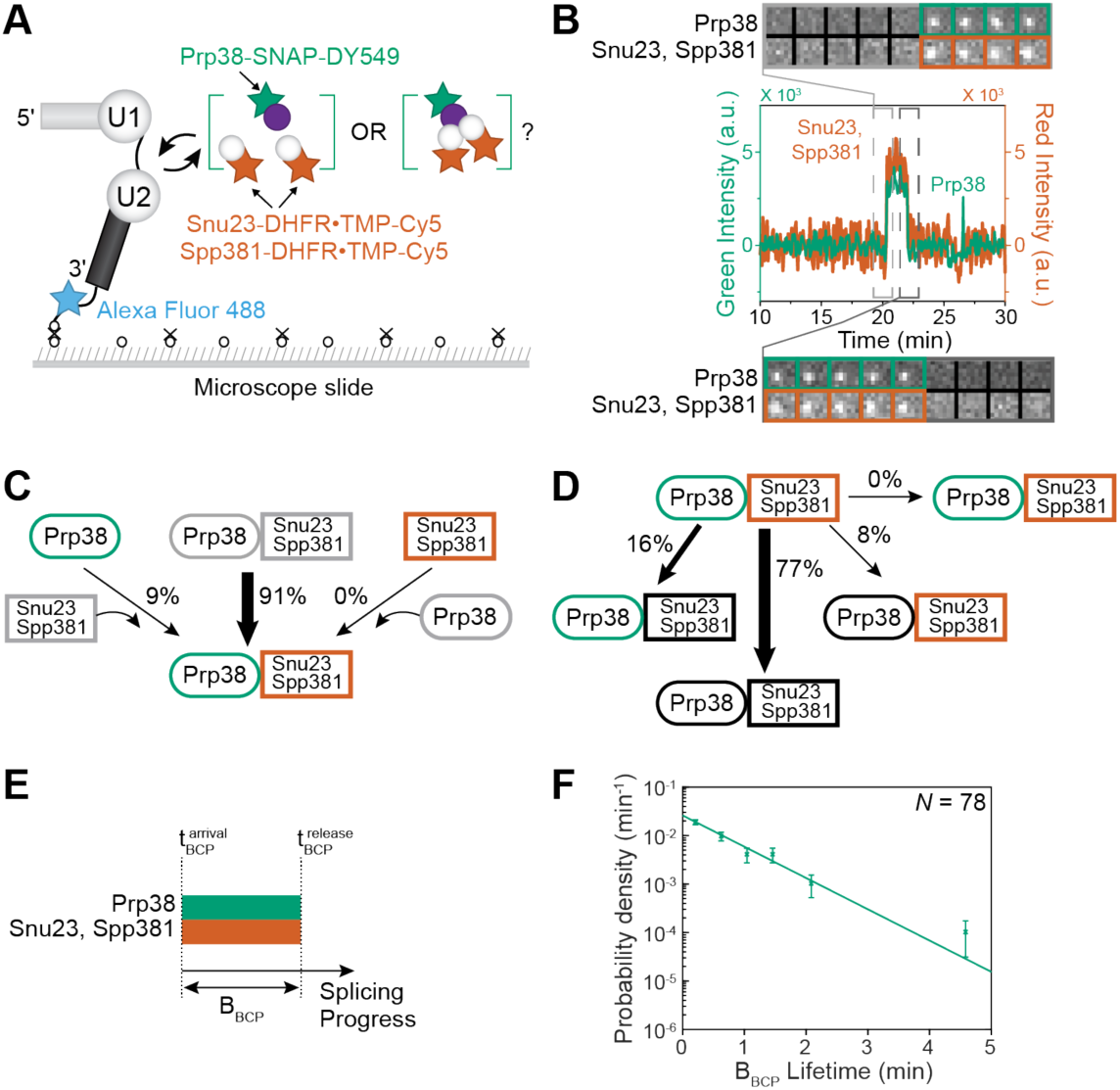
Prp38, Snu23, and Spp381 simultaneously bind to and release from spliceosomes. **(A)** Schematic of a 3-color CoSMoS assay in which Spp381/Snu23 are labeled with Cy5-TMP, Prp38 is labeled with a DY549, and the surface-tethered pre-mRNA is labeled with Alex Fluor 488. **(B)** Representative time record segment showing peaks in fluorescence intensity corresponding to colocalization of Snu23 and Spp381 (red) with Prp38 (green) on the same pre-mRNA molecule. The dashed rectangles mark examples of the simultaneous appearance and disappearance of Snu23, Spp381 and Prp38 spots; galleries show consecutive images taken from the indicated part of the recording showing that both spot appearance and disappearance are simultaneous. **(C)** Routes for recruitment of either the Prp38 or Spp381/Snu23 fluorescent spots at 2 mM ATP for *N*=112 pairs of overlapping events. Green and red shapes represent observation of fluorescence from the corresponding DY549 or Cy5 fluorophores on Prp38 or Snu23, Spp381, respectively. Grey colors represent the absence of fluorescence. Percentages represent the fraction of complexes in which fluorescence appeared by the indicated pathway; more prevalent pathways are emphasized with thicker arrows. (**D**) Routes for loss of either the Prp38 or Snu23, Spp381 fluorescent spots at 2 mM ATP for *N*=102 pairs of overlapping events that showed simultaneous recruitment. Black shapes represent the disappearance of fluorescence. (**E**) Schematic depicting how the lifetime of the B_BCP_ complex is defined relative to the recruitment and release times of BCP-constiuent proteins. (**F**) Probability density histograms of B_BCP_ complex lifetimes obtained from the subset of events (*N*) showing both simultaneous arrival and departure of Prp38, Snu23, and Spp381. The green line represents a fit of the distribution to an equation containing a single exponential term that yielded the parameters reported in **Supplementary Table S3** (τ−_1_ ∼41 s). Error bars (±SD) were calculated for each point as described in the Methods.

We first identified events in which DY549 signals from Prp38 and Cy5 signals from Spp381 and Snu23 were colocalized on the immobilized RNAs since during spliceosome activation Prp38, Snu23, and Spp381 should all be simultaneously present (**Fig. 2B, Supplementary S4**). For these events, the binding patterns were then classified based on the relative recruitment and release orders for Prp38 and Snu23/Spp381 (**Supplementary Fig. S5**). For example, we determined if the DY549 signal appeared or disappeared before, after, or simultaneously with the Cy5 signal for each colocalized pair. We found that most of the event pairs (91%; 102/112) showed simultaneous arrival of Prp38-SNAP and Snu23/Spp381-DHFR signals (**Fig. 2C**). Among this subset, 77% showed simultaneous release of all three proteins (**Fig. 2D**). While it is possible that the proteins individually bind and dissociate too quickly one after the other for our measurements to detect, our data is most consistent with structural and gel filtration data (4,5,14,18) supporting formation of a Prp38/Snu23/Spp381 ternary complex in the absence of a spliceosome (*i.e.*, the BCP). The BCP binds to and releases from the spliceosome as a single unit.

We additionally used the measured dwell times for the colocalized event pairs to determine the lifetime of the BCP-containing spliceosome (the B_BCP_ complex, **Fig. 2E**). The distribution of lifetime measurements could be described by a function containing a single exponential term, resulting in a characteristic lifetime of ∼41 s for B_BCP_ (**Fig. 2F, Supplementary Table S3**). Our determined lifetime of the B_BCP_ species is unlikely to be limited by photobleaching since simultaneous bleaching of the DY549 and Cy5 signals should be rare and we were readily able to measure lifetimes >10-fold larger for the SNAP-labeled proteins when ATP was limiting (**Fig. 1D, Supplementary Table S2**). Since splicing *in vitro* occurs over tens of minutes, dissolution of the B_BCP_ complex occurs rapidly enough to support splicing and likely involves just a single, rate-limiting kinetic step.

### The BCP Joins the Spliceosome after tri-snRNP Recruitment under Splicing Conditions

We then studied when the BCP subcomplex is recruited to the spliceosome and when it is released during activation relative to the U4 snRNP. The U4 snRNP is a core component of the tri-snRNP and thus reports on tri-snRNP binding to the pre-mRNA. Release of U4 snRNP is a key event during the B to B^act^ complex transition since it permits formation of the U2/U6 di-snRNA catalytic site. We carried out 3-color CoSMoS assays using extracts containing SNAP-tagged Snu23 labeled with SNAP-DY549 fluorophore and DHFR-tagged U4 snRNP proteins (Prp3 and Prp4) labeled with Cy5-TMP fluorophores along with surface-immobilized Alexa Fluor 488-labeled pre-mRNAs (**Fig. 3A, Supplementary S6A**). We first identified all paired binding events in which fluorescent signals from the U4 snRNP proteins were simultaneously present on the pre-mRNA with signals from Snu23. We found that most of the event pairs (89%; 31/35 events) showed sequential arrival. The U4 snRNP protein signals appeared first followed by appearance of Snu23, suggesting that the BCP is not recruited as part of the tri-snRNP (**Fig. 3B, C, Supplementary S7A**). Consistently, the same recruitment pattern was also found when extracts containing SNAP-tagged Spp381 or Prp38 were used along with DHFR-tagged U4 snRNP proteins (**Supplementary Figs. S6B-C, S7B-E**). Together these results indicate that the BCP is recruited to the spliceosome after the tri-snRNP associates.

**Figure 3.**
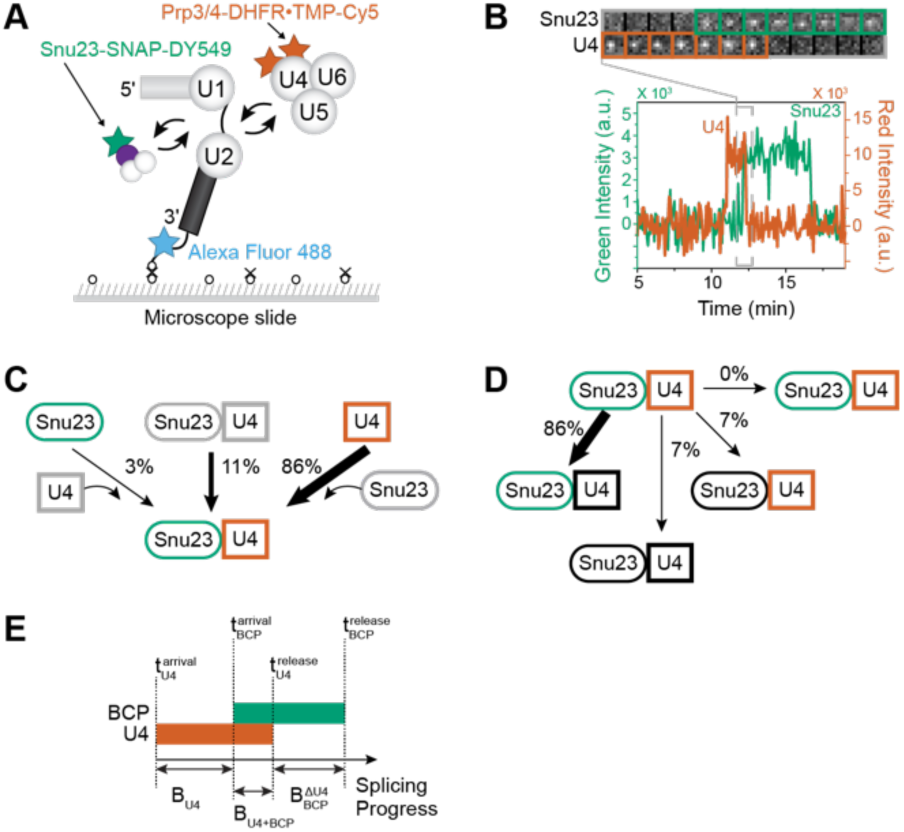
BCP association occurs after tri-snRNP binding and persists after U4 release. (**A**) Schematic of a 3-color CoSMoS assay in which U4 snRNP proteins (Prp3, Prp4) are labeled with Cy5-TMP, the BCP is labeled with a DY549 (here, Snu23), and the surface-tethered pre-mRNA is labeled with Alex Fluor 488. (**B**) Representative time record segment showing peaks in fluorescence intensity corresponding to colocalization of the U4 snRNP (red) with Snu23 (green) on the same pre-mRNA molecule. The dashed rectangle marks an example of ordered recruitment of the tri-snRNP and BCP, followed by U4 release. The image galleries show consecutive images taken from the indicated part of the recording showing that both spot appearances and disappearances are not simultaneous. (**C**) Routes for the appearance of Snu23 and U4 fluorescent spots at 2 mM ATP for *N*=35 pairs of overlapping events. Red and green shapes represent observation of fluorescence from the corresponding DY549 or Cy5 fluorophores on Snu23 or U4, respectively. Percentages represent the fraction of complexes in which fluorescence appeared by the indicated pathway; more prevalent pathways are emphasized with thicker arrows. (**D**) Routes for loss of either Snu23 or U4 fluorescent spots at 2 mM ATP for *N*=30 pairs of overlapping events in which the U4 spot appearance preceded arrival of Snu23. (**E**) Schematic depicting how the lifetimes of the 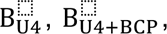, and 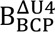 complexes are defined relative to the recruitment and release times of constituent factors. Corresonding data for Prp38 and Spp381 are included in **Supplementary Figs. S6-S10**.

We further analyzed the order of BCP and U4 snRNP release for pairs of colocalized events. In the case of the BCP protein Snu23, the majority of event pairs (86%) showed loss of signals from the U4 snRNP proteins occurring before loss of signals from Snu23 (**Fig. 3D, Supplementary S7A**). This was also the predominant pattern for signal loss from Spp381 and Prp38 when WCEs containing those labeled proteins were used (**Supplementary Figs. S7B-C, F-G**). This result is consistent with ordered release of the U4 snRNP and then the BCP from the spliceosome as well as a previously determined structure of the human pre-B^act-1^ activation intermediate that contains the human BCP but not the U4 snRNP (5). Thus, the yeast spliceosome appears to share with humans conserved pathways for both recruitment and release of BCP proteins during activation.

The single molecule data also indicate the existence of at least three types of spliceosome B complexes: 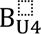 in which the tri-snRNP has associated with the pre-mRNA but the BCP has not yet joined, B_U4+BCP_ in which the BCP has bound but U4 has not been released, and 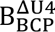 in which the U4 snRNP has been released and the BCP has not yet dissociated (**Fig 3E**). In this nomenclature, we use the subscript to identify components of the B complex spliceosome and the superscript to signify loss of a particular subunit.

We were able to determine the characteristic lifetimes of each complex to be ∼57s, ∼13s, and ∼101s for the B_U4_, B_U4+BCP_,and 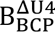 complexes, respectively (**Fig S8, Supplementary Table S3**). These data indicate that under splicing conditions, the BCP and U4 snRNP only transiently co-exist in the same spliceosome complex before U4 snRNP release is completed.

### Snu66 is Released with the U4 snRNP and Before the BCP

We next tested if another protein released during the B to B^act^ transition but not part of the BCP also dissociates subsequent to U4 snRNP release. We carried out three-color CoSMoS assays using WCEs containing Snu66-SNAP labeled with SNAP-DY549 fluorophore and doubly DHFR-tagged U4 snRNP proteins (Prp3 and Prp4) labeled with Cy5-TMP fluorophores (**Fig. 4A**). As before, we identified colocalized pairs of events and then studied the orders in which the signals appeared or were lost. At 2 mM ATP, the predominant pathway for Snu66 association with the pre-mRNA is coincident with U4 snRNP proteins (80%; 77/96 event pairs, **Fig. 4B, C; Supplementary Fig. S9**). Among the subset in which Snu66 and U4 snRNP protein signals appeared simultaneously, the majority also showed simultaneous loss of the signals (72%, 55/77 event pairs, **Fig. 4B, D**). While we cannot exclude the possibility that Snu66 binding or release occurs very swiftly (< 4 s) relative to tri-snRNP binding and U4 dissociation, our data are most consistent with Snu66 being recruited to spliceosomes as part of the tri-snRNP and being released along with the U4 snRNP. Release of Snu66 along with U4 snRNP proteins could be due to direct interaction between Snu66 and Prp3, as has been observed in human tri-snRNPs (29).

**Figure 4.**
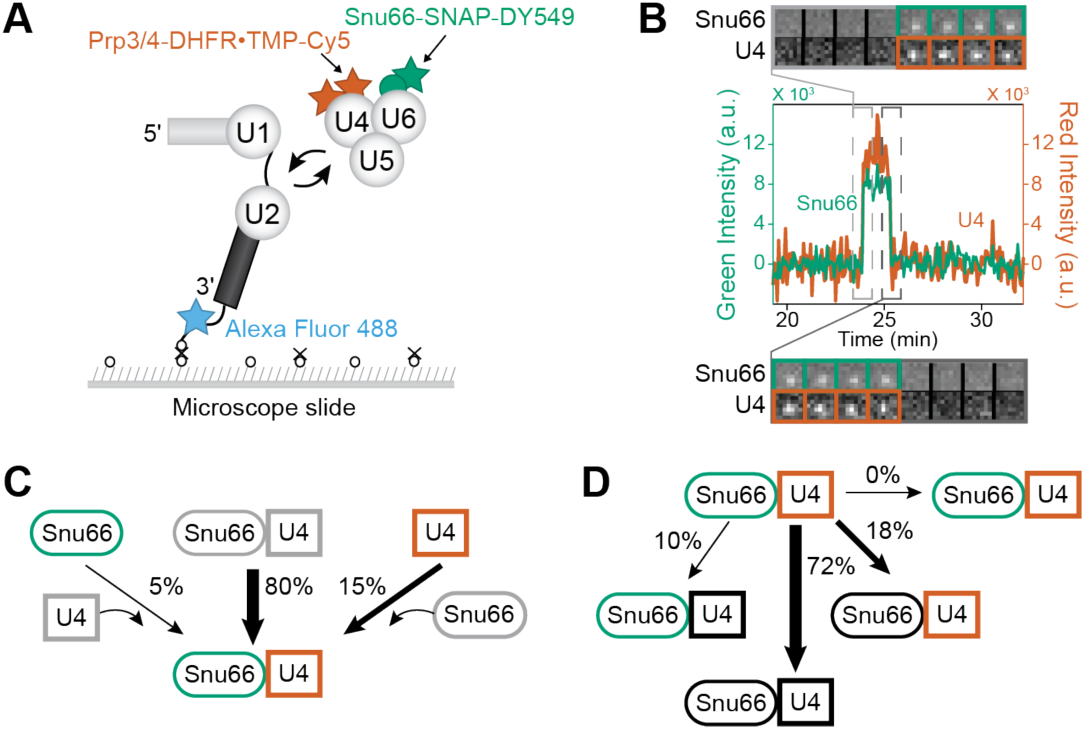
Snu66 is released along with the U4 snRNP during activation. **(A**) Schematic of a 3-color CoSMoS assay in which U4 snRNP proteins (Prp3, Prp4) are labeled with Cy5-TMP, Snu66 is labeled with a DY549, and the surface-tethered pre-mRNA is labeled with Alex Fluor 488. (**B**) Representative time record segment showing peaks in fluorescence intensity corresponding to colocalization of the U4 snRNP (red) with Snu66 (green) on the same pre-mRNA molecule. The dashed rectangles mark examples of the simultaneous appearance and disappearance of U4 and Snu66 spots; galleries show consecutive images taken from the indicated part of the recording. (**C**) Routes for the appearance of Snu66 and U4 fluorescent spots at 2 mM ATP for *N*=96 pairs of overlapping events. Red and green shapes represent observation of fluorescence from the corresponding DY549 or Cy5 fluorophores on Sun66 or U4, respectively. Percentages represent the fraction of complexes in which fluorescence appeared by the indicated pathway; more prevalent pathways are emphasized with thicker arrows. (**D**) Routes for loss of either Snu66 or U4 fluorescent spots at 2 mM ATP for *N*=77 pairs of overlapping events in which Snu66 and U4 bound the pre-mRNA simultaneously.

The above data show that not all proteins released during the B to B^act^ transition dissociate simultaneously. When combined with the data from **Fig. 3**, we were able to predict that Snu66 and the BCP should be released sequentially. Specifically, since the BCP is released after the U4 snRNP dissociates and Snu66 dissociates along with the U4 snRNP, then the BCP should also be released after Snu66. To verify this prediction, we carried out a 3-color CoSMoS assay using extracts containing Snu66-SNAP labeled with SNAP-DY549 fluorophore and doubly DHFR-tagged B complex proteins (Spp381 and Snu23) labeled with Cy5-TMP fluorophores (**Supplementary Fig. S10A**). As expected, 84% (66/79) of event pairs showed that Snu66 associates with the pre-mRNA prior to the BCP (**Supplementary Fig. S10B, C**). Among this subset of event pairs, 67% (44/66) showed that BCP is released after Snu66 (**Supplementary Fig. S10D**), consistent with our prediction.

### The BCP is a tri-snRNP Component at Low ATP

The observation that the BCP is associating with the pre-mRNA after tri-snRNP binding was unexpected since the BCP has previously been identified as a tri-snRNP component by mass spectrometry (10,11,15,30) and was present in samples used to determine the cryo-EM structure of the tri-snRNP, although not modeled(31). One difference between those experiments and ours is that we carried out our single molecule assays under conditions that permit spliceosome activation and splicing (2 mM ATP). Under these conditions, the yeast tri-snRNP is unstable. Consequently, structural and analytical experiments were instead carried out in the absence of added ATP (12,31). We wondered if the presence or absence of ATP changes BCP association with the tri-snRNP. To test this, we repeated our three-color CoSMoS assays using labeled BCP and U4 snRNP but under conditions that permit spliceosome assembly but inhibit activation (0.05 mM ATP) (**Fig. 5A**).

**Figure 5.**
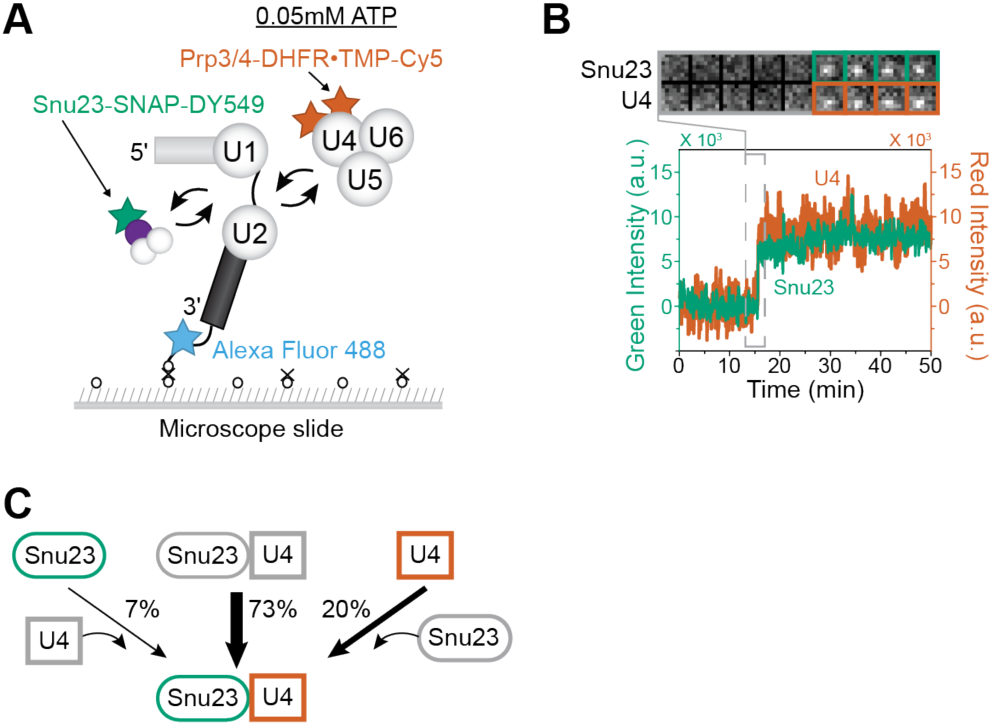
BCP association with the tri-snRNP is ATP-dependent. (**A**) Schematic of a 3-color CoSMoS assay in which U4 snRNP proteins (Prp3, Prp4) are labeled with Cy5-TMP, the BCP is labeled with a DY549 via Snu23, and the surface-tethered pre-mRNA is labeled with Alex Fluor 488. A low concentration of ATP (0.05 mM) is used to inhibit spliceosome activation. (**B**) Segment of a representative time record showing peaks in fluorescence intensity corresponding to colocalization of U4 (red) and Snu23 (green) with the same pre-mRNA molecule. The dashed rectangle marks an example of the simultaneous appearance of U4 and Snu23 spots; galleries show consecutive images taken from the indicated part of the recording showing. (**C**) Routes for the appearance of Snu23 and U4 fluorescent spots at 0.05 mM ATP for *N*=93 pairs of overlapping events. Red and green shapes represent observation of fluorescence from the corresponding DY549 or Cy5 fluorophores on Snu23 or U4, respectively. Percentages represent the fraction of complexes in which fluorescence appeared by the indicated pathway; more prevalent pathways are emphasized with thicker arrows. Corresponding data for the BCP labeled on either Spp381 and Snu23 are included in **Supplementary Fig. S11**.

At 0.05 mM ATP, Snu23 no longer predominantly associated with the pre-mRNA separately from the tri-snRNP. Instead, the most common pathway was simultaneous binding of Snu23 and U4 snRNP proteins (73%; 68/93 event pairs, **Fig. 5B, C, Supplementary S11A**). Simultaneous binding with U4 snRNP proteins was also observed for Spp381, Prp38 and Snu66 when these proteins were labeled instead of Snu23 (**Supplementary Figs. S11B-G**). This observation likely explains why the BCP and Snu66 showed similar binding kinetics in two-color CoSMoS assays at 0.05 mM but not 2 mM ATP (**Fig. 1D, E**). These results indicate that under conditions that stall spliceosome activation, the BCP is recruited to the pre-mRNA as a stable component of the tri-snRNP and remains stably bound to these spliceosome complexes.

### The BCP is Released after NTC Recruitment

Finally, we wished to order release of the BCP relative to recruitment of the NTC. We previously determined that the NTC is predominantly recruited after U4 snRNP release (20). Since the BCP is also released after U4 departure, we wondered if this occurs coincident with, before, or after NTC recruitment. To answer this question, we carried out 3-color CoSMoS assays using WCE containing SNAP-tagged BCP (via Snu23 labeling with SNAP-DY549 fluorophore) and two DHFR tags on NTC components (Syf1 and Cef1) labeled with Cy5-TMP fluorophores (**Fig. 6A**).

**Figure 6.**
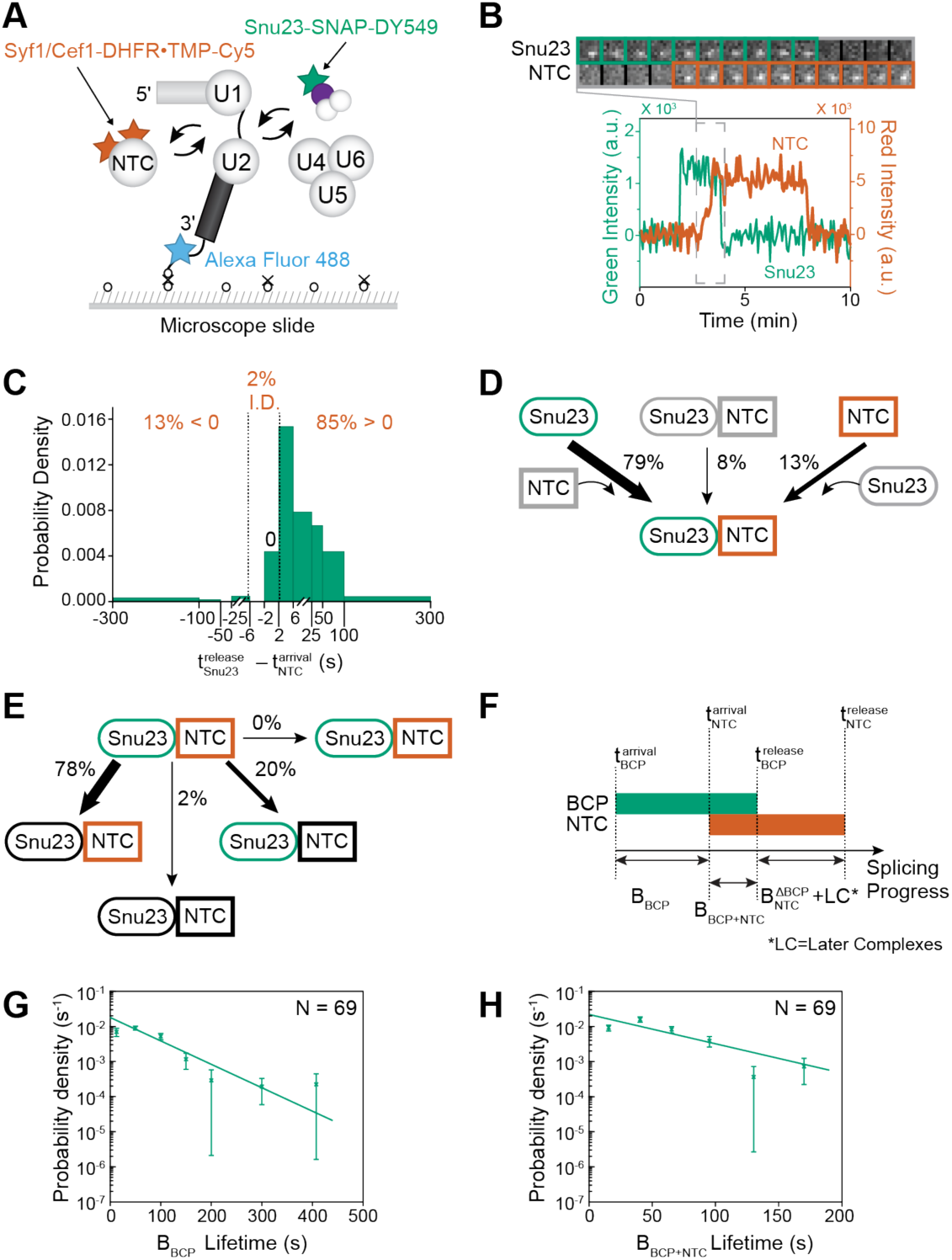
Three-color CoSMoS observation of NTC and Snu23 binding dynamics during activation. (**A**) Schematic of a 3-color CoSMoS assay in which NTC proteins (Syf1, Cef1) are labeled with Cy5-TMP, the BCP is labeled with a DY549 via Snu23, and the surface-tethered pre-mRNA is labeled with Alex Fluor 488. (**B**) Segment of a representative time record showing peaks in fluorescence intensity corresponding to colocalization of NTC (red) and Snu23 (green) with the same pre-mRNA molecule. The light grey dashed rectangle marks an example of the ordered recruitment of the NTC followed by the release of Snu23; galleries show consecutive images taken from that part of the recording. (**C**) Probability density histogram of measured delays between the times of NTC arrivals and Snu23 release events. Most often (85% of *N*= 117 total events), the NTC arrived before release of the Snu23 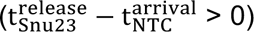. In 2% of cases, the exact order of events was indeterminate (I.D.) due to simultaneous loss of both signals or both signals remaining at the conclusion of the experiment. (**D**) Routes for the appearance of Snu23 and NTC fluorescent spots at 2 mM ATP for *N*=116 pairs of the subset of overlapping events. Red and green shapes represent observation of fluorescence from the corresponding DY549 or Cy5 fluorophores on Snu23 or the NTC, respectively; grey shapes represent the absence of fluorescence. Percentages represent the fraction of complexes in which fluorescence appeared by the indicated pathway; more prevalent pathways are emphasized with thicker arrows. (**E**) Routes for loss of either Snu23 or NTC fluorescent spots at 2 mM ATP for *N*=92 pairs of overlapping events in which the Snu23 spot appearance preceded arrival of the NTC. (**F**) Schematic depicting how the lifetimes of the 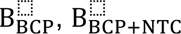, and 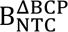 complexes are defined relative to the recruitment and release times of constituent factors. (**G-H**) Probability density histograms of 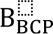 (Panel G, 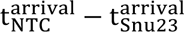) and 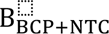 (Panel H, 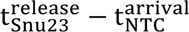) obtained from the subset of events (*N*) showing ordered arrival of Snu23 and then NTC spots followed by ordered loss of the Snu23 and then NTC signals. Lines represent fits of the lifetime distributions with equations containing single exponential terms that yielded the parameters reported in **Table S3** (τ_1_ ∼66 and ∼53 s for 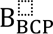 and 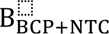, respectively). Error bars (±SD) were calculated for each point as described in the Methods.

We identified pairs of BCP and NTC binding events closest in time to one another with a requirement that binding events for both components should be at least 2 frames in duration (≥ 8 s) to eliminate sampling interactions of the NTC (19). For each pair, we then subtracted the time of NTC binding 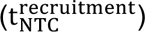 from the time of BCP signal loss 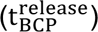 to yield a distribution of 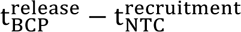 values. In this distribution, positive values would indicate that BCP is released after NTC recruitment while negative values would indicate that BCP release occurs before NTC recruitment. The distribution of 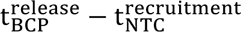 values for the selected pairs of events showed that 85% had positive values (97/114 events), suggesting that the NTC is recruited while the BCP is still present and that the BCP is released after the NTC binds (**Figs. 6B-E; Supplementary Figure S12**). In comparison, the same analysis carried out for extracts containing SNAP-tagged Snu66 and doubly DHFR-tagged NTC proteins showed predominantly negative values, consistent with loss of Snu66 preceding NTC recruitment (**Supplementary Fig. S13**).

The single molecule data also indicate the existence of several intermediate complexes: B_BCP_in which the BCP has associated with the pre-mRNA but the NTC has not yet joined, B_BCP+NTC_ in which the NTC and BCP are both simultaneously present, and 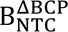 and later complexes in which the BCP has released but the NTC remains (**Fig 6F**). The characteristic lifetimes of the B_BCP_and B_BCP+NTC_ complexes are relatively short (∼1 min; **Fig. 6G, H, Supplementary Table S3**). The total lifetime of the Snu23-SNAP-labeled BCP in these assays was 9-fold shorter than that for the fluorophore measured under the same conditions but attached to an immobilized SNAP protein (∼110 s vs. ∼995 s) (19). This supports the conclusion that the measured lifetimes are indicative of binding and release of the BCP from splicing complexes rather than being primarily due to photobleaching. The B_BCP+NTC_ intermediate has not yet been described for yeast spliceosomes and likely possesses a similar composition and structure to the human pre-B^act-1^ complex (5).

## DISCUSSION

Integrating previous work from our group (19,20) with single molecule data described here, we can derive a kinetic scheme for spliceosome activation *in vitro* (**Fig. 7A**). In our scheme, the tri-snRNP, lacking the BCP, joins the spliceosome A complex containing U1 and U2 to form the pre-B complex. After ∼57s, the BCP joins as a single subunit to form the B_U4+Lsm+BCP_ complex.

**Figure 7.**
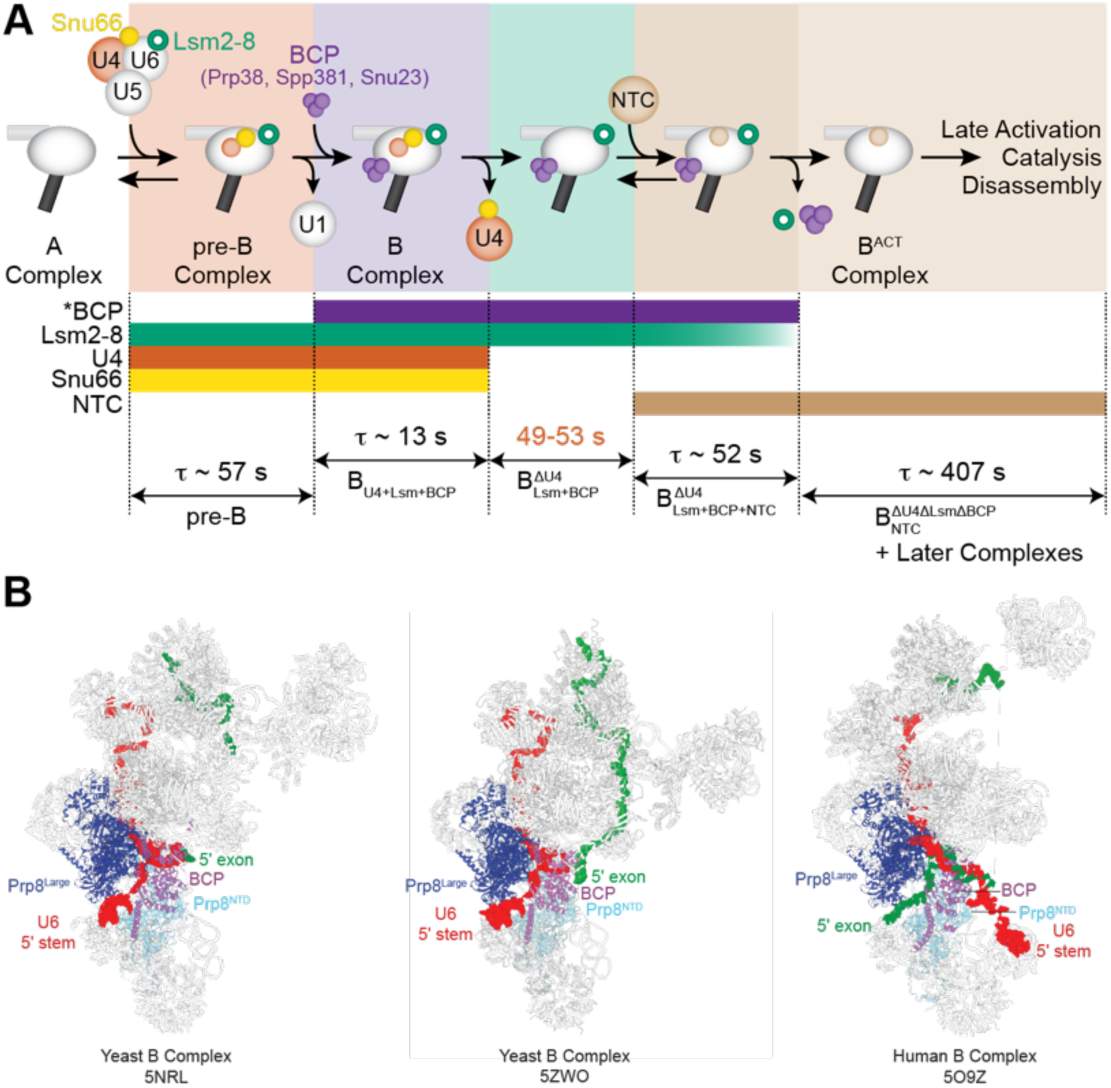
A Kinetic Scheme for Spliceosome Activation. (**A**) In this kinetic scheme, a spliceosome complex containing an intact tri-snRNP (as evidenced by presence of the U4 snRNP and Lsm2-8 complexes) and the BCP 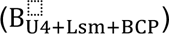 forms very transiently (τ ∼13 s) before the U4 snRNP is released to form the 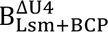 complex. After ∼50 s, the NTC joins to form the 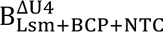 spliceosome. The Lsm2-8 and BCP complexes are then released in an undetermined order after another ∼50 s. The lifetime of the 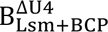 complex (red) was not directly measured in these experiments but can be deduced from our studies in combination with previously reported experiments that measured the delay between U4 release and NTC recruitment (19,20). (**B**) Structural alignment of yeast (left, middle) and human (right) B complex spliceosomes highlights differences in positioning of the U6 snRNA 5’ stem region (red) and 5’ exon (green) of the pre-mRNA substrates in yeast and human spliceosomes. Spliceosomes were aligned based on the Prp8 N-terminal domain (Prp8^NTD^) using PyMol and visualized using PyMol and ChimeraX software.

Based on cryo-EM structures of yeast and human spliceosomes, the BCP associates near the U6 snRNA/5’ SS duplex at the same site vacated by Prp28 (5,32,33). Thus, the ∼57s time interval reports on both BCP association and transfer of the 5’ SS from the U1 snRNA to the U6 snRNA by Prp28. We have not yet analyzed U1 snRNP dynamics during activation, but it is likely that it also dissociates during this time.

The lifetime of B_U4+Lsm+BCP_ is very short, ∼13 s, before the U4 snRNP is lost along with at least a subset of tri-snRNP specific factors including Snu66. This forms a spliceosome complex lacking U4 and Snu66 but containing the Lsm2-8 and BCP complexes 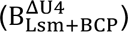. This complex then persists for 49-53 s, before the NTC associates to form the 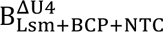 complex. 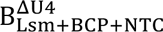 in turn persists for another ∼52 s before dissociation of the BCP and Lsm2-8 complexes to form the 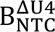 complex, which likely has a similar composition and structure to the B^act^ spliceosome (34,35). We do not yet know the order of BCP and Lsm2-8 complex departure, as we have not yet been able to assay this particular step (not shown). Once B^act^ is formed, spliceosomes usually persist for ∼407 s before loss of the NTC due to either successful splicing or disassembly of stalled complexes.

Currently, no structural data is available for yeast 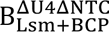 or 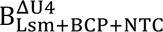 complexes. However, it is likely that they may be structurally analogous to the human pre-B^act-1^ and pre-B^act-2^ complexes, respectively, with the exception that yeast likely recruit the NTC as a single subunit while the Lsm complex remains bound (5,19). Importantly, our model is consistent with evolutionary conservation of BCP recruitment and release between yeast and humans: the BCP arrives independent of the tri-snRNP to B complex and is released after U4 snRNP dissociation and arrival of at least a partial complement of NTC proteins (5). Since our work indicates that yeast 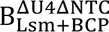 or 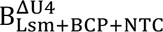 complexes exist only transiently, obtaining cryo-EM structural data for these may benefit from either synchronizing spliceosome activation followed by rapid freezing of the samples at short time intervals or use of small molecules or splicing factor mutations that result in accumulation of these states.

### BCP Association with the tri-snRNP is ATP Dependent

It has previously been proposed that BCP components are part of the tri-snRNP and recruited to the spliceosome as part of this assembly. This was largely based on mass spectrometry data and biochemical assays of complexes isolated under no or limiting ATP conditions (10–13,15,30). Conversely, data from the human splicing machinery indicates that the corresponding human proteins are recruited during activation after tri-snRNP binding (33,36). Our CoSMoS data resolve these observations. Under conditions that permit activation and splicing, the BCP is recruited to the spliceosome after incorporation of the tri-snRNP, in agreement with a model proposed by the Shi laboratory based on cryo-EM structures of endogenous yeast pre-B and B complexes (32). This mechanism simplifies models for yeast spliceosome activation by eliminating the need to reconcile BCP occupancy near the 5’ SS with Prp28 activity. Stepwise recruitment of the tri-snRNP and BCP to spliceosomes would allow Prp28 to transfer the 5’ SS from U1 to U6 and dissociate from the spliceosome prior to BCP binding. The BCP can then be positioned to stabilize interactions and conformational changes needed for subsequent steps in activation (5,7).

However, limiting ATP results in the BCP being recruited to the yeast spliceosome as part of the tri-snRNP. Thus, the BCP is an ATP-dependent component of the yeast tri-snRNP, and this explains its presence in complexes isolated under low ATP conditions. Whether or not the BCP could similarly be recruited to human tri-snRNPs in the absence of spliceosome formation is uncertain. Human tri-snRNPs appear to be stabilized in an unactivated conformation with Brr2 located far from its U4 snRNP substrate and with stably-bound Prp28 (36). These factors may preclude premature BCP association.

### Premature Association of the BCP May Result in Unproductive Spliceosomes

We believe that yeast tri-snRNPs formed at low ATP and containing the BCP are unlikely to be competent for splicing. Previous single molecule data showed that the majority (68%) of spliceosomes formed at 0.05 mM ATP were disassembled at 2 mM ATP (20). In addition, tri-snRNPs purified in the absence of ATP and containing the BCP disassemble when ATP is added (31). This would support the hypothesis that many of the BCP-containing tri-snRNP complexes that bind to the pre-mRNA are not able to carry out subsequent steps and represent non-functional species. It should be noted; however, that we have not directly assayed the splicing competency of the BCP-associated tri-snRNP.

Formation of non-functional yeast spliceosome complexes at low ATP is also supported by analysis of cryo-EM data for yeast and human B complexes. Importantly, the yeast B complex spliceosomes were assembled at low, 0.05 mM ATP (5NRL) or purified from yeast in the absence of ATP (5ZWO) whereas the human B complex was isolated at high ATP (2 mM) using a low Mg^2+^ concentration (5O9Z) (4,14,32). When comparing these structures, it is obvious that the 5’ stem of the U6 snRNA and 5’ exon of the pre-mRNA have exchanged locations (**Fig. 7B**). In catalytic spliceosomes from both yeast and humans (B* or C complexes that form after the B complex), the 5’ exon is accommodated in an “exon channel” formed by Prp8. In the human B spliceosome, the 5’ exon is already located in the exon channel and the 5’ SS is base paired with the U6 snRNA ACAGAGA sequence proximal to the channel (**Fig. 7B**, right). These interactions are stabilized by the human BCP. However, in the published yeast B spliceosome complexes the 5’ SS has not yet base paired with the U6 ACAGAGA sequence and is instead pairing with an adjacent upstream element (the ACAGAGA stem). Additionally, the 5’ exon is located outside the channel while the U6 5’ stem is located inside, stabilized by the BCP (**Fig. 7B**, left and middle). Previously, the differences in 5’ SS/U6 base-pairing and around the exon channel were interpreted as divergent activation mechanisms taken by yeast and human splicing machinery with yeast requiring additional steps to correctly pair U6 and position the 5’ exon (37).

We propose that yeast B complex structures formed at limiting ATP represent off-pathway or non-functional intermediates due to pre-mature association of the BCP with the tri-snRNP. The BCP may structurally lock the U6 5’ stem in the exon channel of the B complex as well as block correct transfer of the 5’ SS to U6 by preventing Prp28 binding. This results in formation of spliceosomes not competent for carrying out the splicing reaction and that are substrates for disassembly at higher concentrations of ATP. It is interesting to note that insertion of the U6 snRNA 5’ stem into the exon channel was also observed in structures of endogenous spliceosomes purified from lysates (**Fig. 7B**, middle) (32). This suggests that formation of this structure may be biologically relevant and not specific to just one pre-mRNA substrate.

Whether or not BCP-dependent formation of non-functional tri-snRNP or spliceosome complexes at low ATP is used to regulate splicing *in vivo* is unknown. It is possible that switching of the tri-snRNP between catalytically competent and incompetent conformations depending on the presence or absence of the BCP could be used to globally regulate cellular splicing in response to ATP levels. In *S. cerevisiae*, cellular ATP concentration is usually maintained at a constant level of ∼4 mM (38). However, under certain conditions such as deletion of the Bas1 transcription factor that regulates purine biosynthesis or in the presence of glycolysis inhibitors such as 2-deoxyglucose, intracellular levels can fall dramatically including to levels that we would predict would cause BCP association with the tri-snRNP (≤ 0.05 mM) (39). Under such conditions, “spliceosome shutdown” caused by formation of BCP-associated tri-snRNPs and non-functional spliceosomes could lead to reduced ribosome synthesis (since the majority of yeast introns are found within ribosomal protein genes (40)) and a redirection of cellular ATP towards other metabolic or cellular needs.

## DATA AVAILABILITY

Single molecule data (microscope video recordings) will be deposited in Zenodo upon manuscript acceptance/publication.

## SUPPLEMENTARY DATA

Supplementary data (Supplemental Figures 1-13 and Supplemental Tables 1-3) accompany this manuscript.

## ACKNOWLEDGEMENTS

We thank Laura Vanderploeg (Dept. of Biochemistry, UW-Madison) for help in creating the figures and illustrations. We also thank Dr. Kathy Senn and Ethan Aubuchon for critical reading of the manuscript.

## Author Contributions

XF and AAH conceptualized the project; XF collected and analyzed the data; XF and AAH wrote the manuscript.

## FUNDING

This work was supported by grants from the National Institutes of Health (R35 GM136261 to AAH).

## CONFLICT OF INTEREST DISCLOSURE

AAH is a member of the scientific advisory board and carrying out sponsored research for Remix Therapeutics.

