## Supplemental Figures and Tables for "Dynamics and Evolutionary Conservation of B Complex Protein Recruitment During Spliceosome Activation"

This supplementary file contains:

Supplementary Figures S1-S13

Supplemetnary Tables S1-S3

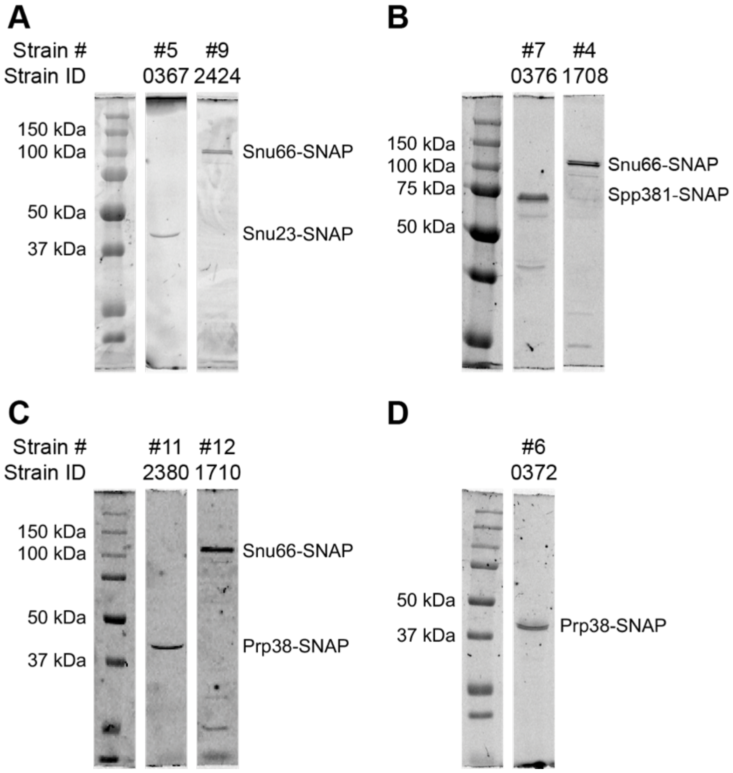


**Supplementary Figure S1. Fluorophore Labeling of Extracts of Various SNAP-tagged Strains.** SNAP-tagged proteins can be specifically labeled in yeast WCE with SNAP-DY549. Shown are fluorescence images of SDS-PAGE gels for confirming successful labeling. The gel images were cropped (vertical white stripes) to remove intervening lanes not relevant to this figure.

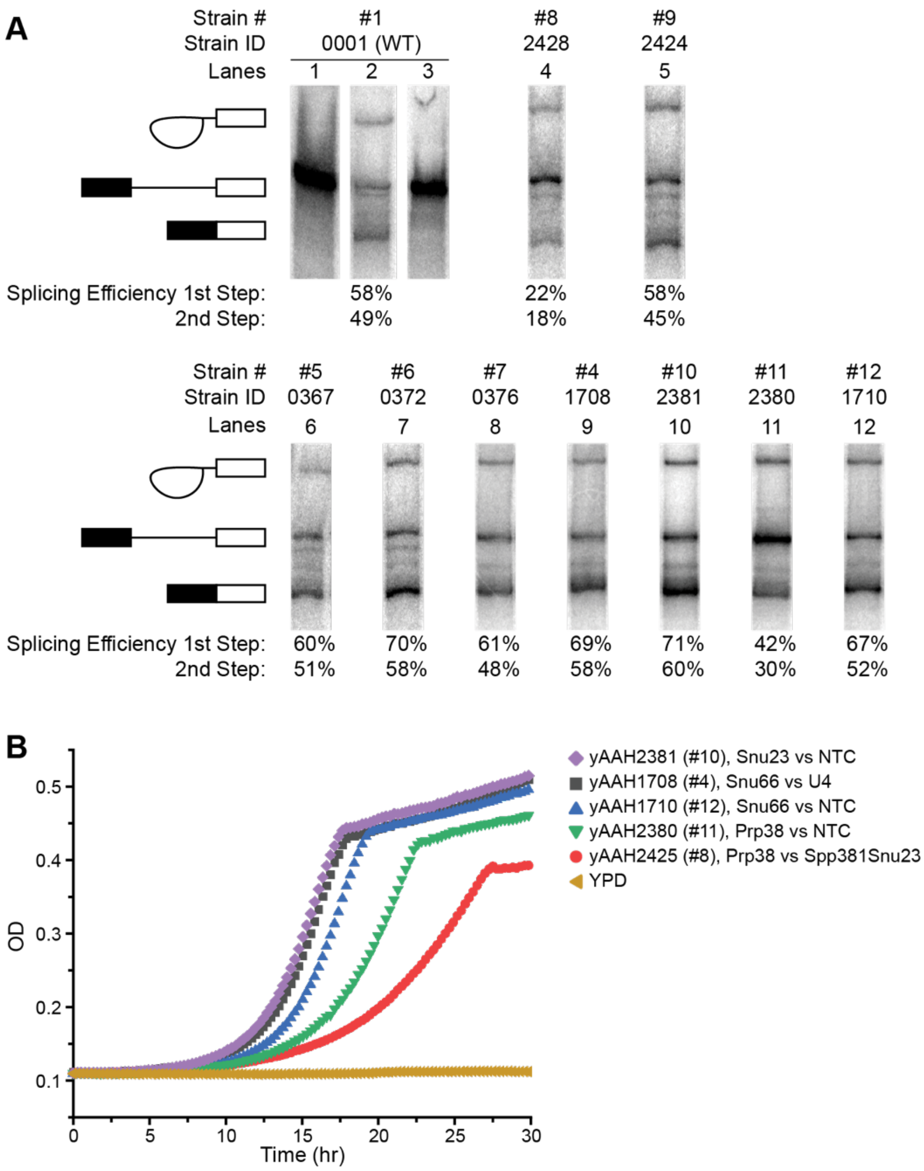


**Figure S2. Splicing Activities of Dy549-Labeled WCE.** (**A**) Representative splicing assays and efficiencies for WCE containing fluorophore-labeled proteins (lanes 4-12) or a WT control (lanes 1-3). In lanes 4-12, ATP was added to a concentration of 2 mM to promote splicing. In lane 3, ATP was added to a concentration of 0.05 mM to permit spliceosome assembly but prevent activation and splicing. Lanes 1 represents a sample at t=0 min while the other lanes represent reactions quenched at 45 min. Cartoons for splicing products are indicated to the left of the gel. The gel images were cropped (vertical white stripes) and combined into a single figure to remove intervening lanes not relevant for this figure and for clarity. (**B**) Yeast growth assay results for the strains shown indicating slower growth of Prp38-tagged strains.

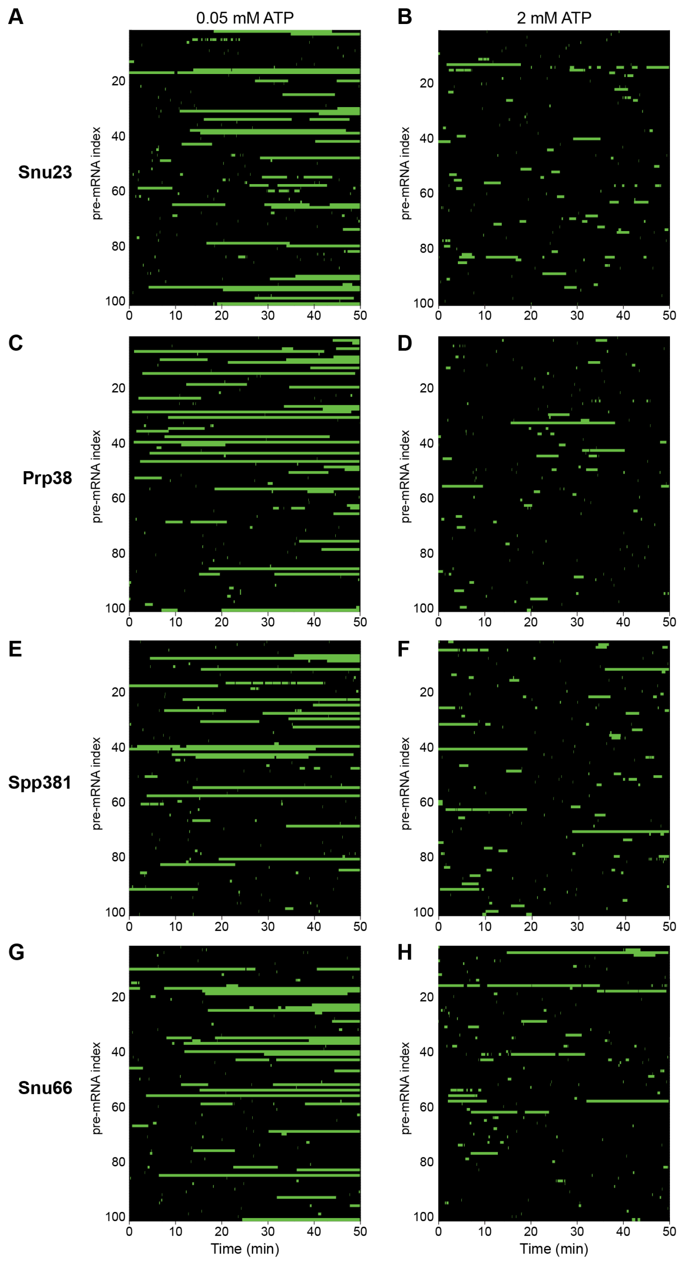


**Figure S3. ATP-Dependent Binding Intervals of BCP Components (Prp38, Snu23, and Spp381) and Snu66 on Single pre-mRNA Molecules.** Shown are binding intervals (rastergrams) of fluorescently labeled proteins on single pre-mRNA molecules under conditions that inhibit (0.05 mM ATP, left columns, long intervals) or permit activation and splicing (2 mM ATP, right columns, short intervals). Within each panel, each row represents association and dissociation of fluorescent molecules on an individual pre-mRNA molecule over 50 min, with green and black bars representing fluorescent and dark states.

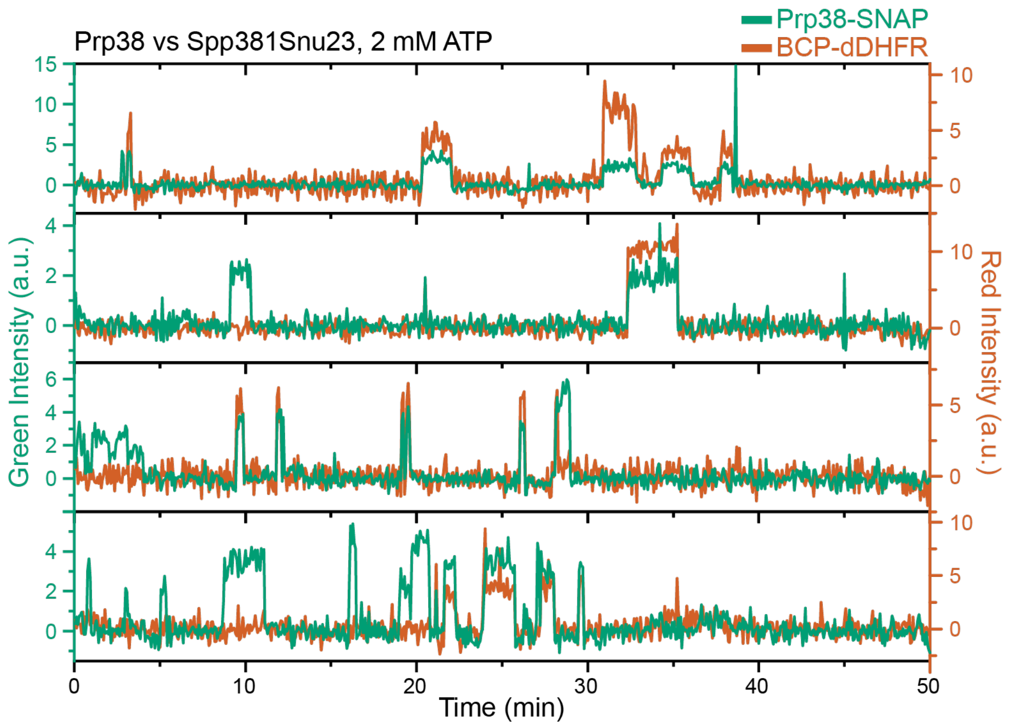


**Figure S4. Additional Sample Fluorescence Trajectories from 3-color CoSMoS Experiments Monitoring Prp38-SNAP and BCP (Snu23, Spp381)-DHFR Proteins.** Shown are super-imposed fluorescence intensities for Prp38-SNAP (green traces) and BCP-DHFR proteins (red traces) at 2 mM ATP.

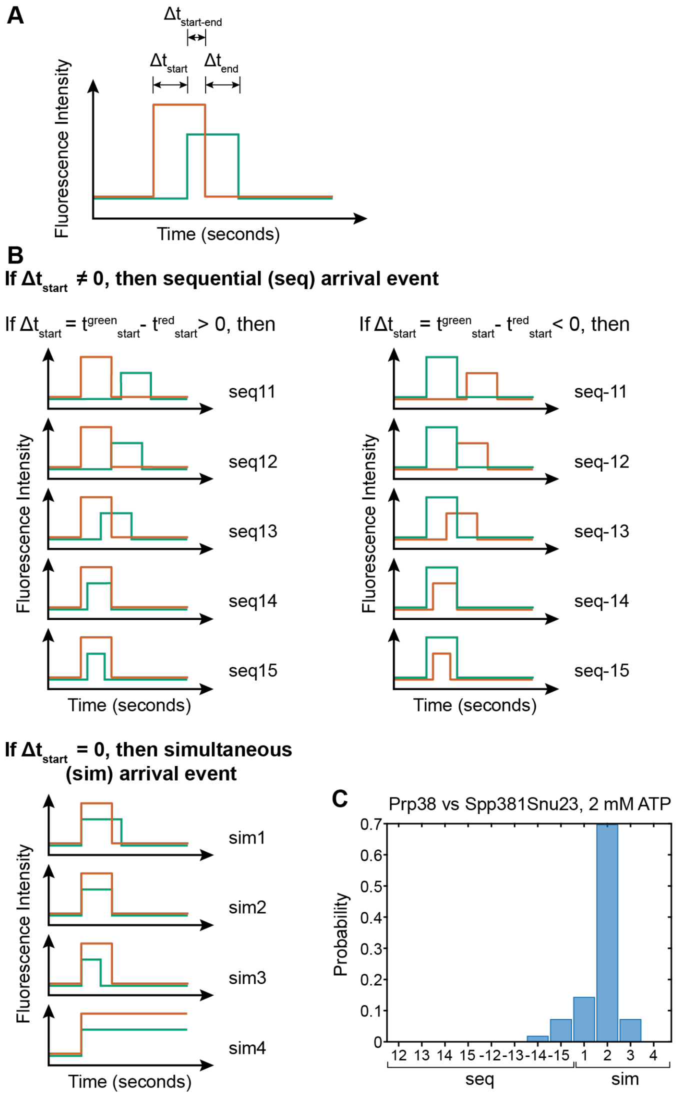


**Figure S5. Scheme for assignment of observed binding and dissociation patterns in CoSMoS assays.** (**A**) For each set of paired events, the beginning and end times of the individual events were recorded. For the analysis shown here, $\Delta t_{start}$ values were first calculated by subtracting the start time of the green event from the start time of the red event. (**B**) $\Delta t_{start}$ values were separated such that all positive values indicated that the red event arrived before the green event (seq11 to 15), negative values indicated that the green event arrived after the red event (seq-11 to -15), and zero values indicated simultaneous arrival (sim1 to 4). Values of $\Delta t_{end}$ and $\Delta t_{start\text{-}end}$ were then determined to subcategorize each arrival class based on the disappearance patterns of the green and red events. (**C**) Distribution of Prp38 and BCP overlapped binding events from 3-color CoSMoS experiments shows the majority are in the sim2 class.

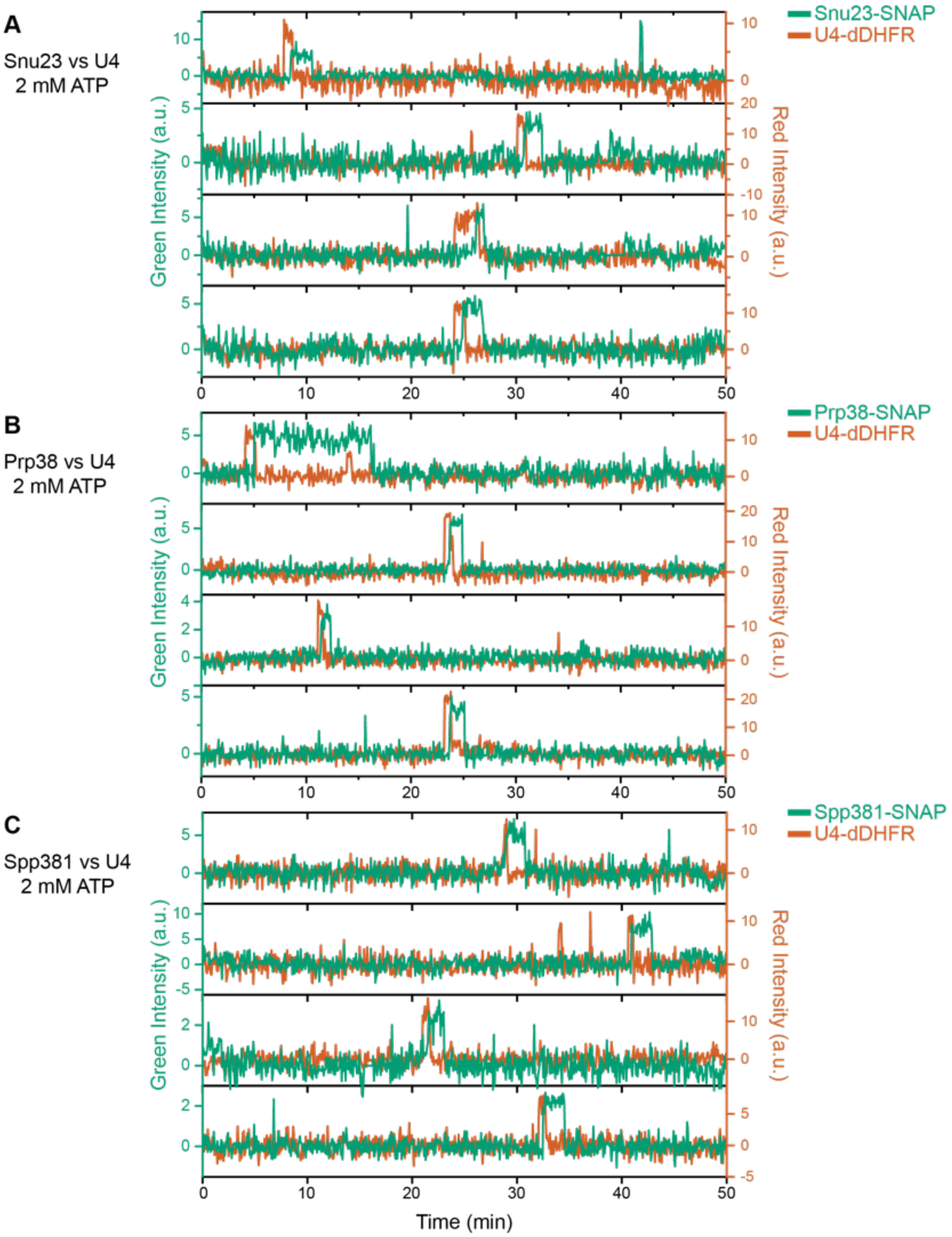


**Figure S6. Additional Sample Fluorescence Trajectories from 3-color CoSMoS Experiments Monitoring BCP-SNAP and U4-DHFR Proteins.** Shown are super-imposed fluorescence intensities for Prp38-SNAP, Snu23-SNAP or Spp381-SNAP (green traces) and BCP-DHFR proteins (red traces) at 2 mM ATP.

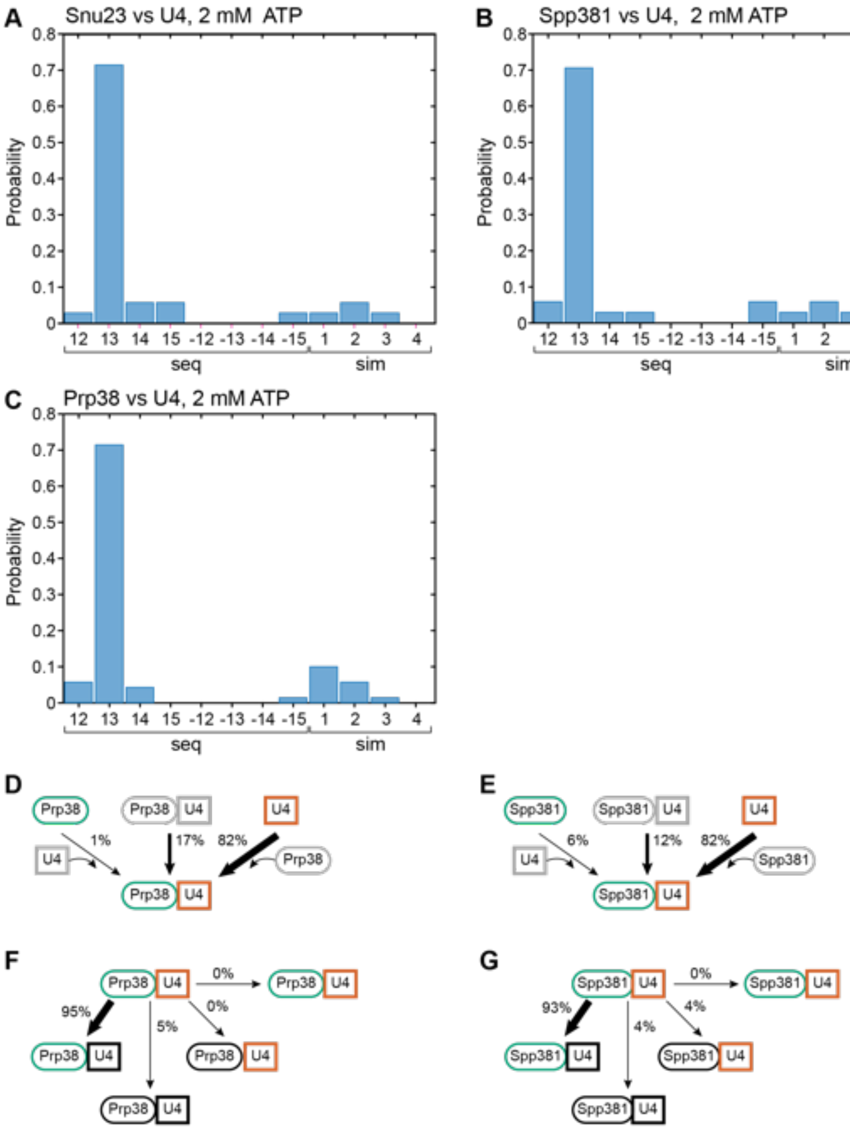


**Figure S7. Additional Analysis on BCP and U4 binding dynamics during activation.** (**A-C**) Distributions of overlapped BCP and U4 binding events from 3-color CoSMoS experiments. (**D-G**) Routes for the appearance of Prp38 (**D**) or Spp381 (**E**) and U4 fluorescent spots at 2 mM ATP for *N*=70 or 34 pairs of overlapping events, respectively. (**F-G**) Routes for loss of Prp38 (**F**) or Spp381 (**G**) and U4 fluorescent spots at 2 mM ATP for *N*=51 or 25 pairs of overlapping events for Prp38 and Spp381, respectively, in which the U4 spot appearance preceded arrival of the BCP component.

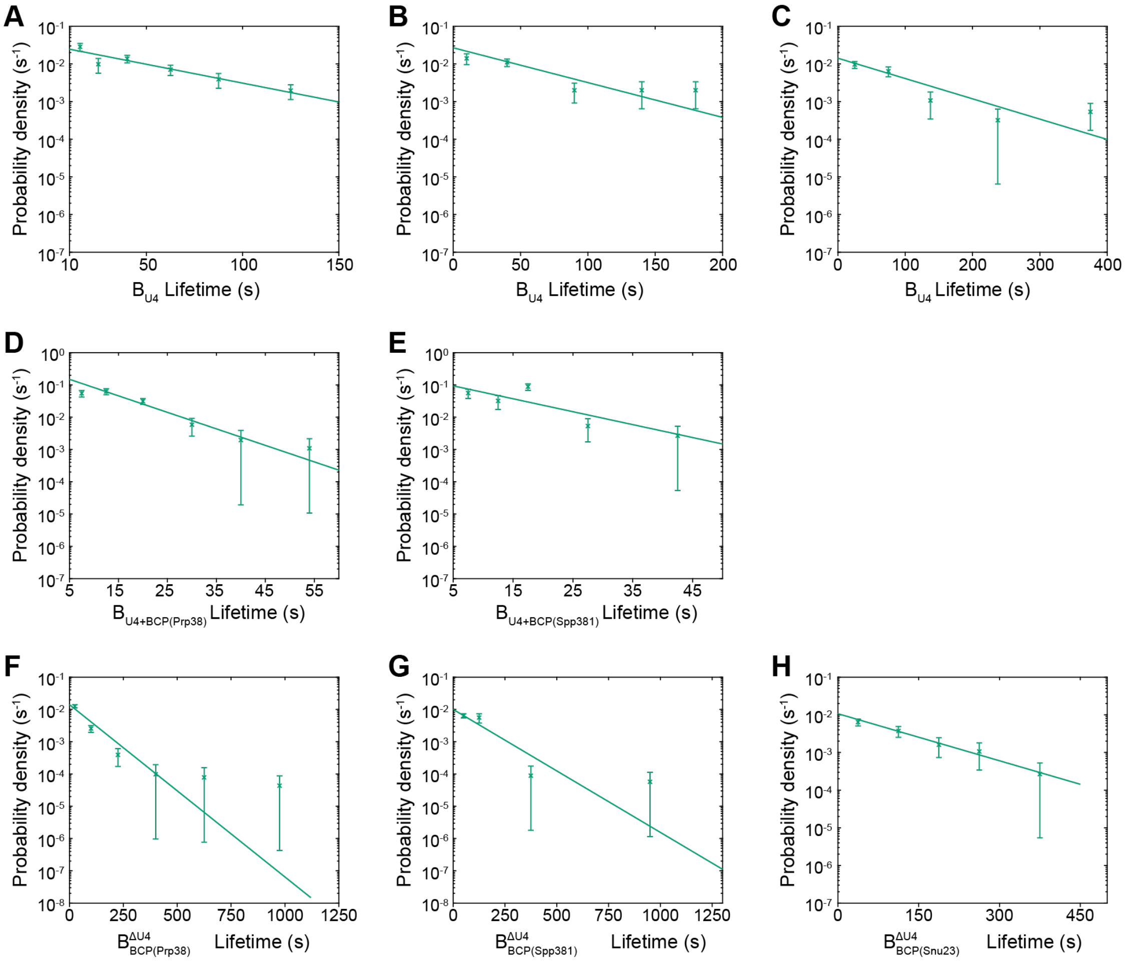


**Figure S8. Probability Density Histograms of Intermediates from Observing BCP and U4 Binding Dynamics.** Probability density histograms of $\mathrm{B}_{\mathrm{U4}}^{\square}$ (panel **A-C**, $\mathrm{t}_{\mathrm{BCP}}^{\mathrm{arrival}} - \mathrm{t}_{\mathrm{U4}}^{\mathrm{arrival}}$), $\mathrm{B}_{\mathrm{U4+BCP}}^{\square}$ (panel **D-E**, $\mathrm{t}_{\mathrm{U4}}^{\mathrm{release}} - \mathrm{t}_{\mathrm{BCP}}^{\mathrm{arrival}}$) and $\mathrm{B}_{\mathrm{BCP}}^{\Delta\mathrm{U4}}$ (panel **F-H**, $\mathrm{t}_{\mathrm{BCP}}^{\mathrm{release}} - \mathrm{t}_{\mathrm{U4}}^{\mathrm{release}}$) obtained from the subset of events showing ordered arrival of U4 and then BCP spots followed by ordered loss of the BCP and then U4 signals. For Panels A-C, these represent $\mathrm{B}_{\mathrm{U4}}^{\square}$ values obtained from extracts with SNAP-labeled Spp381, Prp38, and Snu23, respectively. Lines represent fits of the lifetime distributions with equations containing single exponential terms that yielded the parameters reported in **Supplementary Table S3**. The number of data points included in each fit (*N*) are also reported in **Supplementary Table S3** Error bars were calculated for each point as described in the Methods.

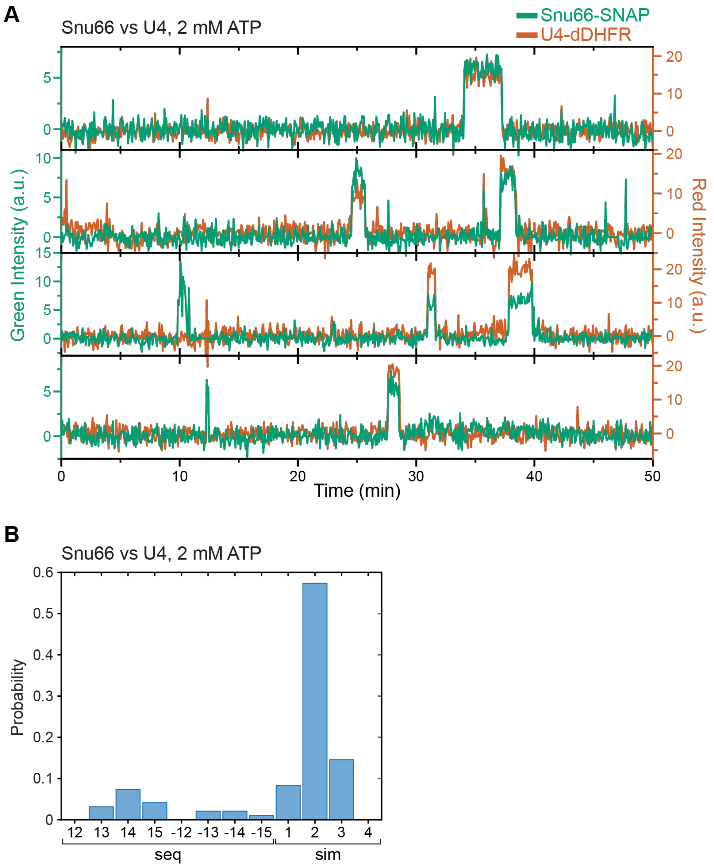


**Figure S9. Additional Trajectories and Analysis from Observing Snu66 and U4 Binding Dynamics. (A)** Shown are super-imposed fluorescence intensities for Snu66-SNAP (green traces) and U4-DHFR proteins (red traces) at 2 mM ATP. (**B**) Distribution of Snu66 and U4 overlapped binding events from 3-color CoSMoS experiments.

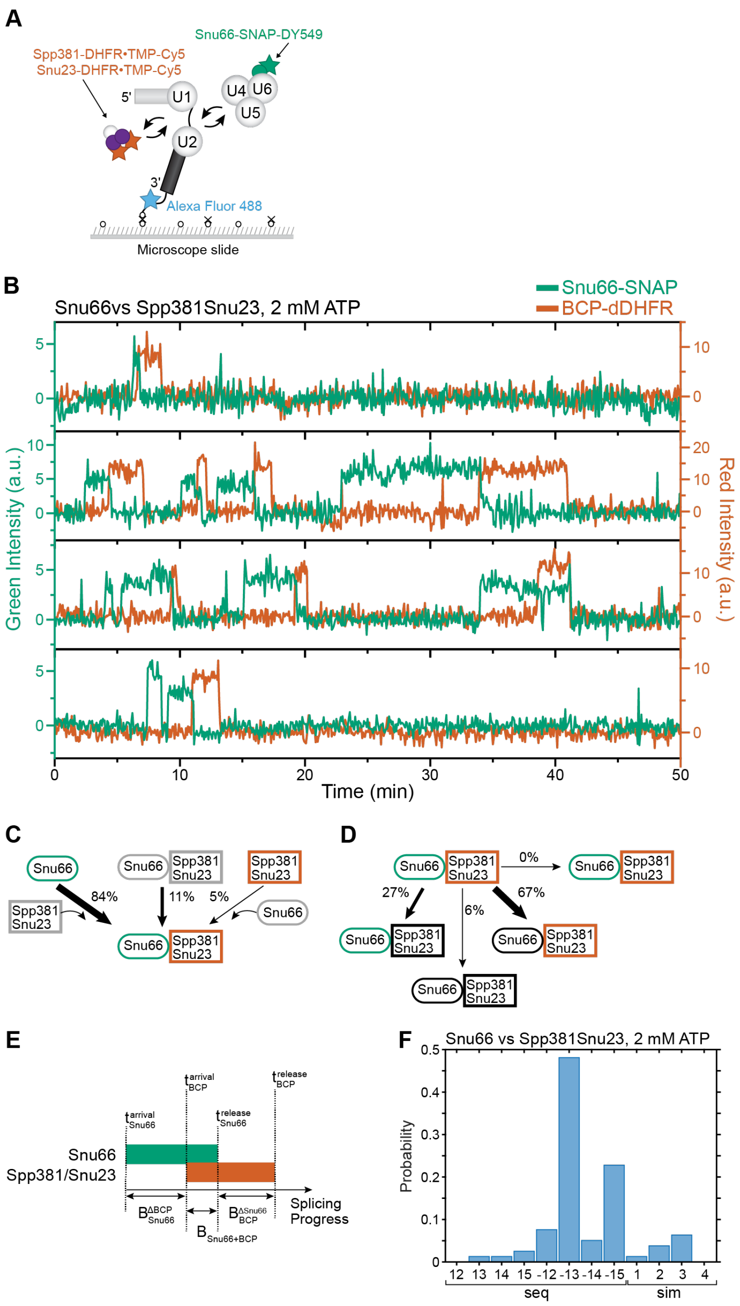


**Figure S10. 3-color CoSMoS observation of Snu66 and BCP binding dynamics during activation.** (**A**) Schematic of a 3-color experiment in which the BCP was labeled with Cy5 fluorophores via Snu23 and Spp381, Snu66 was labeled with a DY549 fluorophore, and the surface-tethered pre-mRNA was labeled with an Alexa Fluor 488 fluorophore. (**B**) Shown are super-imposed fluorescence intensities for Snu66-SNAP (green traces) and BCP-DHFR proteins (red traces) at 2 mM ATP. (**C**) Routes for the appearance of Snu66 and BCP fluorescent spots at 2 mM ATP for $N$=79 pairs of overlapping events. (**D**) Routes for loss of either Snu66 or BCP fluorescent spots at 2 mM ATP for $N$=66 pairs of overlapping events in which the Snu66 spot appearance preceded arrival of BCP. (**E**) Schematic showing the definition for the identified intermediate based on the relative recruitment and release times of Snu66 and BCP proteins. (**F**) Distribution of Snu66 and BCP overlapped binding events from 3-color CoSMoS experiments.

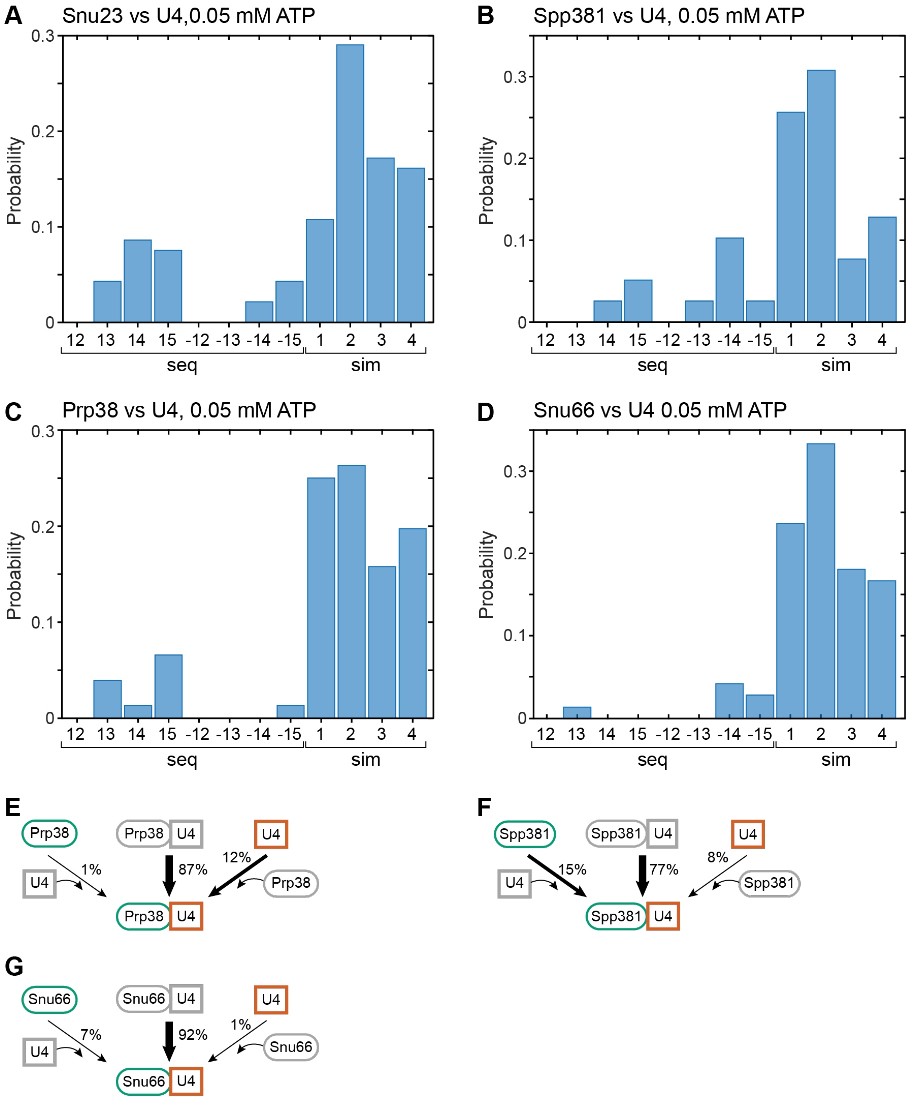


**Figure S11. Additional Analysis on BCP and U4 binding dynamics at low ATP (0.05 mM).** (**A-D**) Distribution of BCP and U4 binding events from 3-color CoSMoS experiments at 0.05 mM ATP. (**E-G**) Routes for the appearance of Prp38 (**E**), Spp381 (**F**) or Snu66 (**G**) and U4 fluorescent spots at 0.05 mM ATP for *N*=76, 39 or 72 pairs of overlapping events, respectively.

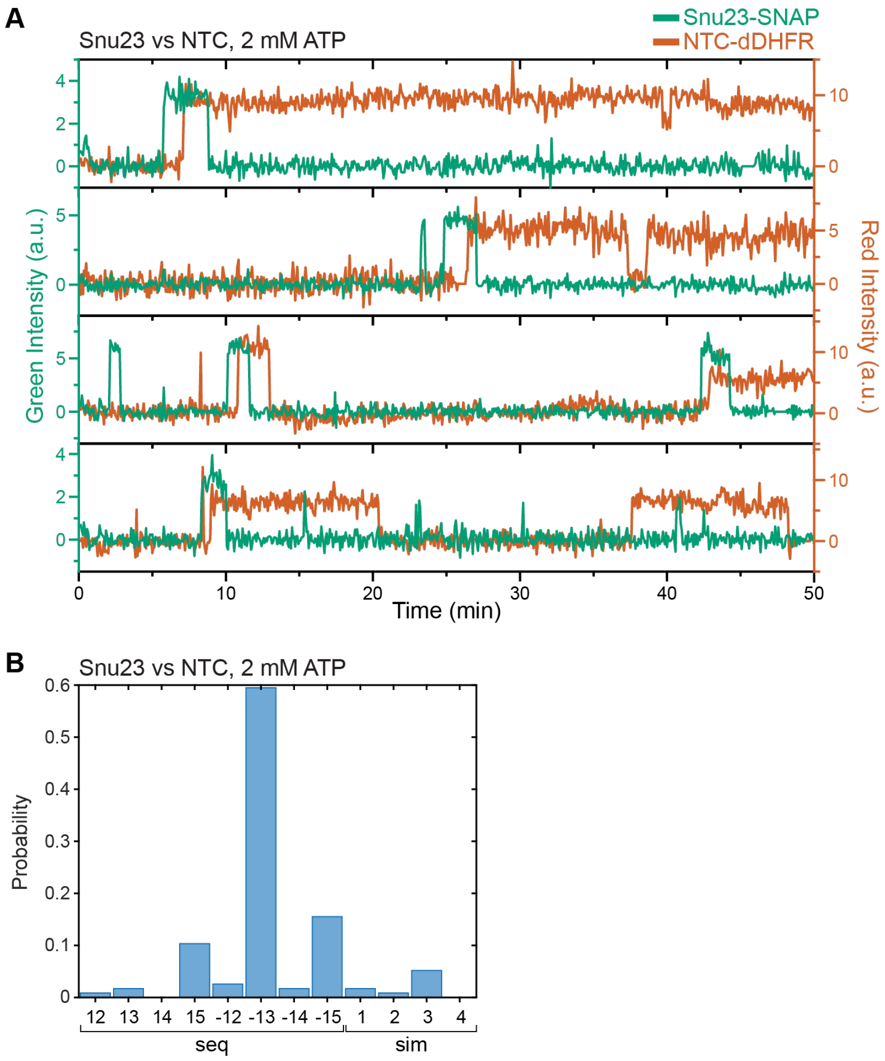


**Figure S12. Additional Trajectories and Analysis from Observing Snu23 and NTC Binding Dynamics.** **(A)** Shown are super-imposed fluorescence intensities for Snu23-SNAP (green traces) and NTC-DHFR proteins (red traces) at 2 mM ATP. (**B**) Distribution of Snu23 and NTC binding events from 3-color CoSMoS experiments.

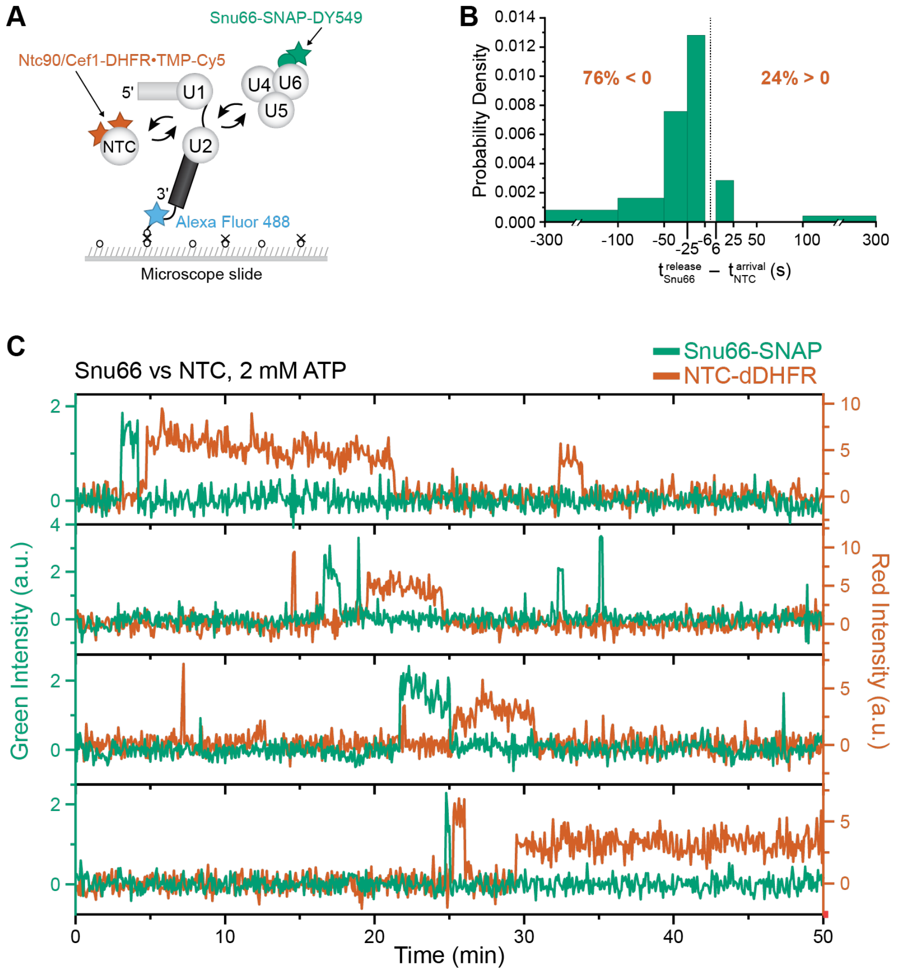


**Figure S13. 3-color CoSMoS observation of Snu66 and NTC binding dynamics during activation.** (**A**) Schematic of a 3-color experiment in which the NTC was labeled with Cy5 fluorophores (via Ntc90(Syf1) and Cef1), Snu66 was labeled with a DY549 fluorophore, and the surface-tethered pre-mRNA was labeled with an Alexa Fluor 488 fluorophore. (**B**) Probability density histogram showing the delay between NTC arrival and Snu66 release. Most often (76% of *N*= 39 total events), the NTC arrived after release of the Snu66 ($t^{release}_{Snu66} - t^{arrival}_{NTC} < 0$). (**C**) Shown are super-imposed fluorescence intensities for Snu66-SNAP (green traces) and NTC-DHFR proteins (red traces) at 2 mM ATP.

**Table S1**

| Strain # | ID | Genotype | Notes |
|---|---|---|---|
| 1 | yAAH0001 | MATa prc1-407 prb1-1122 pep4-3 leu2 trp1 ura3-52 gal2 | BJ2168, parental strain |
| 2 | yAAH0018 | yAAH0001 + cef1::cef1-DHFR-HPH + ntc90::ntc90-DHFR-BLE | contains two DHFR-tagged NTC proteins with hygromycin and phleomycin resistance markers |
| 3 | yAAH0329 | yAAH0001 + prp3::prp3-DHFR-HPH + prp4::prp4-DHFR-BLE | contains two DHFR-tagged U4 snRNP proteins with hygromycin and phleomycin resistance markers |
| 4 | yAAH1708 | yAAH0329 + snu66::snu66-$SNAP_f$-NAT | contains two DHFR-tagged U4 snRNP proteins and fast-SNAP-tagged snu66 with nourseothricin resistance marker |
| 5 | yAAH0367 | yAAH0329 + snu23::snu23-$SNAP_f$-KanMX | contains two DHFR-tagged U4 snRNP proteins and fast-SNAP-tagged snu23 with kanamycin resistance marker |
| 6 | yAAH0372 | yAAH0329 + prp38::prp38-$SNAP_f$-NAT | contains two DHFR-tagged U4 snRNP proteins and fast-SNAP-tagged prp38 with nourseothricin resistance marker |
| 7 | yAAH0376 | yAAH0329 + spp381::spp381-$SNAP_f$-NAT | contains two DHFR-tagged U4 snRNP proteins and fast-SNAP-tagged spp381 with nourseothricin resistance marker |
| 8 | yAAH2425 | yAAH0001 + prp38::prp38-$SNAP_f$-NAT + spp381::spp381-DHFR-HPH + snu23::snu23-DHFR-BLE | contains two DHFR-tagged B complex proteins with hygromycin and phleomycin resistance markers and fast-SNAP-tagged prp38 with nourseothricin resistance marker |
| 9 | yAAH2424 | yAAH0373 + spp381::spp381-DHFR-HPH + snu23::snu23-DHFR-BLE | contains two DHFR-tagged B complex proteins with hygromycin and phleomycin resistance markers and fast-SNAP-tagged snu66 with nourseothricin resistance marker |
| 10 | yAAH2381 | yAAH0018 + snu23::snu23-$SNAP_f$-NAT | contains two DHFR-tagged NTC proteins and fast-SNAP-tagged snu23 with nourseothricin resistance marker |
| 11 | yAAH2380 | yAAH0018 + prp38::prp38-$SNAP_f$-NAT | contains two DHFR-tagged NTC proteins and fast-SNAP-tagged prp38 with nourseothricin resistance marker |
| 12 | yAAH1710 | yAAH0018 + snu66::snu66-$SNAP_f$-NAT | contains two DHFR-tagged NTC proteins and fast-SNAP-tagged snu66 with nourseothricin resistance marker |
| 13 | yAAH0373 | yAAH0001 + snu66::snu66-$SNAP_f$-NAT | contains a fast-SNAP-tagged snu66 with nourseothricin resistance marker |

### Supplementary Table S2

### Fit parameters for binding dynamics of proteins from 2-color CoSMoS experiments

| Tagged protein | Tagged Subcomplex | Strain | [ATP] mM | $a_1$ | $\tau_1$ (s) | $a_2$ | $\tau_2$ (s) | $a_3$ | $\tau_3$ (s) | Number of Events | Related Figure |
|---|---|---|---|---|---|---|---|---|---|---|---|
| Snu66-SNAP | tri-snRNP | yAAH1708 | 2 | 0.89±0.02 | 15.2±0.8 | 0.11±0.02 | 213.0±26.0 | | | 688 | Fig. 1 |
| Snu23-SNAP | BCP | yAAH0367 | 2 | 0.54±0.03 | 6.9±0.5 | 0.46±0.03 | 92.9±8.3 | | | 471 | Fig. 1 |
| Spp381-SNAP | BCP | yAAH0376 | 2 | 0.72±0.02 | 7.9±0.4 | 0.28±0.02 | 97.2±9.2 | | | 684 | Fig. 1 |
| Prp38-SNAP | BCP | yAAH0372 | 2 | 0.53±0.05 | 6.9±0.4 | 0.47±0.03 | 82.7±7.5 | | | 697 | Fig. 1 |
| Snu66-SNAP | tri-snRNP | yAAH1708 | 0.05 | 0.63±0.05 | 7.8±0.7 | 0.23±0.05 | 45.7±14.8 | 0.14±0.07 | 435.8±80.0 | 385 | Fig. 1 |
| Snu23-SNAP | BCP | yAAH0367 | 0.05 | 0.77±0.02 | 8.1±0.5 | 0.07±0.03 | 77.4±49.6 | 0.16±0.04 | 640.0±132.2 | 514 | Fig. 1 |
| Spp381-SNAP | BCP | yAAH0376 | 0.05 | 0.74±0.03 | 7.6±0.6 | 0.17±0.03 | 61.9±21.6 | 0.09±0.04 | 908.3±645.1 | 356 | Fig. 1 |
| Prp38-SNAP | BCP | yAAH0372 | 0.05 | 0.79±0.02 | 8.6±0.6 | 0.21±0.02 | 616.7±87.1 | | | 466 | Fig. 1 |

**Table S3. Fitted Kinetic Parameters from 3-color CoSMoS experiments**

| Complex | Strain | [ATP] mM | $a_1$ | $\tau_1$ (s) | $a_2$ | $\tau_2$ (s) | $a_3$ | $\tau_3$ (s) | Number of Events | Related Figure |
|---|---|---|---|---|---|---|---|---|---|---|
| $\mathrm{B}_{\mathrm{BCP}}$ | yAAH2425 | 2 | NA | 40.6±5.8 | NA | NA | NA | NA | 78 | Fig. 2F |
| $\mathrm{B}_{\mathrm{U4}}$ | yAAH0376 | 2 | NA | 47.0±22.8 | NA | NA | NA | NA | 25 | Fig. S8B |
| $\mathrm{B}_{\mathrm{U4+BCP(Spp381)}}$ | yAAH0376 | 2 | NA | 10.8±2.5 | NA | NA | NA | NA | 25 | Fig. S8E |
| $\mathrm{B}_{\mathrm{BCP(Spp381)}}^{\Delta \mathrm{U4}}$ | yAAH0376 | 2 | NA | 114.0±47.8 | NA | NA | NA | NA | 25 | Fig. S8G |
| $\mathrm{B}_{\mathrm{U4}}$ | yAAH0367 | 2 | NA | 80.5±29.1 | NA | NA | NA | NA | 25 | Fig. S8C |
| $\mathrm{B}_{\mathrm{U4+BCP(Snu23)}}$ | yAAH0367 | 2 | NA | 18.2 | NA | NA | NA | NA | 25 | *See Note |
| $\mathrm{B}_{\mathrm{BCP(Snu23)}}^{\Delta \mathrm{U4}}$ | yAAH0367 | 2 | NA | 108.7±31.2 | NA | NA | NA | NA | 25 | Fig. S8H |
| $\mathrm{B}_{\mathrm{U4}}$ | yAAH0362 | 2 | NA | 43.4±9.9 | NA | NA | NA | NA | 51 | Fig. S8A |
| $\mathrm{B}_{\mathrm{U4+BCP(Prp38)}}$ | yAAH0362 | 2 | NA | 8.5±1.4 | NA | NA | NA | NA | 51 | Fig. S8D |
| $\mathrm{B}_{\mathrm{BCP(Prp38)}}^{\Delta \mathrm{U4}}$ | yAAH0362 | 2 | NA | 81.4±24.2 | NA | NA | NA | NA | 51 | Fig. S8F |
| $\mathrm{B}_{\mathrm{BCP}}$ | yAAH2381 | 2 | NA | 65.5±8.8 | NA | NA | NA | NA | 69 | Fig. 6G |
| $\mathrm{B}_{\mathrm{BCP+NTC}}$ | yAAH2381 | 2 | NA | 52.7±7.2 | NA | NA | NA | NA | 69 | Fig. 6H |
| $\mathrm{B}_{\mathrm{NTC}}^{\Delta \mathrm{BCP}}$ | yAAH2381 | 2 | NA | 409.0±47.9** | NA | NA | NA | NA | 69 | Fig. 6 |

*The characteristic lifetime of $\mathrm{B}_{\mathrm{BCP(Snu23)+U4}}$ could not be fitted by exponential-based functions and was estimated by averaging the measured dwell times.

**The long lifetime of the NTC after release of the BCP is likely impacted by photobleaching and this parameter should be considered an estimate. However, it is consistent with prior measurements of NTC lifetimes (*1*, *20*).
